# Attenuated Anticipation of Social and Monetary Rewards in Autism Spectrum Disorders

**DOI:** 10.1101/2020.07.06.186650

**Authors:** Sarah Baumeister, Carolin Moessnang, Nico Bast, Sarah Hohmann, Julian Tillmann, David Goyard, Tony Charman, Sara Ambrosino, Simon Baron-Cohen, Christian Beckmann, Sven Bölte, Thomas Bourgeron, Annika Rausch, Daisy Crawley, Flavio Dell’Acqua, Guillaume Dumas, Sarah Durston, Christine Ecker, Dorothea L. Floris, Vincent Frouin, Hannah Hayward, Rosemary Holt, Mark H. Johnson, Emily J.H. Jones, Meng-Chuan Lai, Michael V. Lombardo, Luke Mason, Marianne Oldehinkel, Tony Persico, Antonia San José Cáceres, Thomas Wolfers, Will Spooren, Eva Loth, Declan G. M. Murphy, Jan K. Buitelaar, Heike Tost, Andreas Meyer-Lindenberg, Tobias Banaschewski, Daniel Brandeis, the AIMS-2-TRIALS group

## Abstract

**Background:** Reward processing has been proposed to underpin atypical social behavior, a core feature of autism spectrum disorder (ASD). However, previous neuroimaging studies have yielded inconsistent results regarding the specificity of atypicalities for social rewards in ASD. Utilizing a large sample, we aimed to assess altered reward processing in response to reward type (social, monetary) and reward phase (anticipation, delivery) in ASD.

**Methods:** Functional magnetic resonance imaging during social and monetary reward anticipation and delivery was performed in 212 individuals with ASD (7.6-30.5 years) and 181 typically developing (TD) participants (7.6-30.8 years).

**Results:** Across social and monetary reward anticipation, whole-brain analyses (p<0.05, family-wise error-corrected) showed hypoactivation of the right ventral striatum (VS) in ASD. Further, region of interest (ROI) analysis across both reward types yielded hypoactivation in ASD in both the left and right VS. Across delivery of social and monetary reward, hyperactivation of the VS in individuals with ASD did not survive correction for multiple comparisons. Reward type by diagnostic group interactions, and a dimensional analysis of autism trait scores were not significant during anticipation or delivery. Levels of attention-deficit/hyperactivity disorder (ADHD) symptoms did not affect reward processing in ASD.

**Conclusions:** Our results do not support current theories linking atypical social interaction in ASD to specific alterations in processing of social rewards. Instead, they point towards a generalized hypoactivity of VS in ASD during anticipation of both social and monetary rewards. We suggest that this indicates attenuated subjective reward value in ASD independent of social content and ADHD symptoms.

## Introdcution

Altered reward processing has been proposed to underlie the challenges that individuals with autism spectrum disorder (ASD) face in social interactions. The social motivation hypothesis postulates that individuals with ASD from early in development do not perceive social stimuli as rewarding as typically developing (TD) individuals, which may impact the development of social learning and social skills (1).

Neurobiological evidence in favor of the social motivation hypothesis is however mixed. To assess atypical motivation, reward processing is commonly investigated during the anticipation of a potential reward (“wanting”), the delivery of the reward (“liking”) or during both phases. Further, different types of rewards can be assessed, with non-social (usually monetary) rewards being most commonly investigated across psychiatric conditions, while social rewards have been postulated to be specifically impacted in ASD as detailed in the social motivation hypothesis. Supporting the concept of atypical social reward processing in ASD, one study showed reduced activation in the ventral striatum (VS) (2), a key region for reward processing comprising the nucleus accumbens and caudate head (3), compared to control participants when receiving social rewards. A similar effect was observed in another study in more dorsal parts of the striatum (4). However, other studies did not find functional striatal differences between ASD and TD individuals for social rewards during delivery (5, 6) or anticipation (4, 5). Similarly mixed results exist for non-social rewards: while some previous studies report VS hypoactivation in individuals with ASD when receiving monetary rewards (7-9), this has not been found (5) or only at uncorrected thresholds (10) elsewhere. Results for the anticipation of monetary rewards are also inconsistent with some studies suggesting VS hypoactivation in ASD participants (5, 9, 11), while another showed no difference between ASD and TD (4). Some of the inconsistency of previous findings is likely due to the heterogeneity of ASD itself (12), but also to the relatively small sample sizes examined (ranging between 13 and 39 individuals per group). A recent meta-analysis has partly addressed the latter issue by summarizing the current literature (13). Comparing individuals with ASD to TD, the authors reported striatal hypoactivation during social as well as non-social rewards in ASD. However, results differed between anticipation and delivery phases. They report hypoactivation of the left caudate during anticipation of social rewards, and hyperactivation during the anticipation of non-social rewards. In contrast, during reward delivery, striatal (left nucleus accumbens and caudate) hyperactivation to social rewards and right caudate hypoactivation to non-social rewards were observed in ASD. These findings suggest opposing atypicality patterns for social and monetary reward types between reward phases and do not imply typical non-social reward processing in ASD. Across the seven studies assessing social reward processing, caudate hypoactivation was linked (albeit only at trend-level) to severity of autistic traits as measured with the Social Responsiveness Scale (SRS). This meta-analysis was an important first step to provide a more comprehensiveinsight into atypical reward processing in ASD. However, the number of included studies is still small (e.g. only three studies allowed for the differentiation between reward phases for social reward) and should thus be regarded as exploratory. Further, task designs were heterogenous, which might have increased variability in brain responses and distorted task-specific effects. Finally, while some studies included in the meta-analysis administered social and non-social reward conditions in the same sample, some only assessed one type of reward, limiting direct comparability.

Another challenge is the fact that ASD and attention-deficit/hyperactivity disorder (ADHD) frequently co-occur (14) and atypical reward processing for monetary rewards is often reported in individuals with ADHD (15). However, ADHD comorbidity or symptoms have not been examined in the majority of studies exploring reward processing deficits in ASD (for exceptions, see 6, 16).

Hence, the brain functional mechanisms underpinning reward processing alterations in ASD remain unclear. We therefore investigated reward-related brain responses in a large, well-powered sample of individuals with ASD. The Longitudinal European Autism Project (LEAP; (17)) provides a deeply phenotyped dataset of children, adolescents and adults with and without ASD who performed a social and a monetary reward task. The task was chosen based on its ability to reliably elicit VS reward signaling (18) and allows for the analysis of both reward anticipation and reward delivery phases. We comprehensively assessed differences in reward signaling based on clinical diagnosis as well as dimensional autistic traits. Based on the recent meta-analysis (13), compared to TD, we hypothesized that neurofunctional responses in the VS would show a pattern of increased activity in ASD during monetary, and reduced activity during social reward anticipation – and the opposing pattern during reward delivery. We expected to observe this pattern in categorical case-control comparisons as well as in dimensional analyses based on autism traits. Further, based on previous findings (16), we hypothesized an additive effect of ADHD comorbidity, with reward processing being most severely altered in autistic individuals with elevated ADHD symptoms.

## Methods and Materials

### Experimental procedure

#### Sample

In the LEAP study, 437 individuals with ASD and 300 typically developing individuals, aged between 6 and 30 years, underwent comprehensive clinical, cognitive, and MRI assessment at one of six study sites: Institute of Psychiatry, Psychology and Neuroscience, King’s College London, United Kingdom (KCL); Autism Research Centre, University of Cambridge, United Kingdom (UCAM); Radboud University Nijmegen Medical Centre, the Netherlands (RUNMC); University Medical Centre Utrecht, the Netherlands (UMCU); Central Institute of Mental Health, Mannheim, Germany (CIMH); University Campus Bio-Medico of Rome, Italy (UCBM) (17). The study was approved by the local ethical committees of participating centers and written informed consent was obtained from all participants or their legal guardians (for participants <18 years). For further details about the study design we refer to Loth et al. (17), and for a comprehensive clinical characterization of the LEAP cohort we refer to Charman et al. (19). For this study, the final sample consisted of n=213 ASD and n=181 TD participants (see table 1). Standard operating and quality control procedures leading to the final sample are detailed in the supplemental material.

**Table 1:**
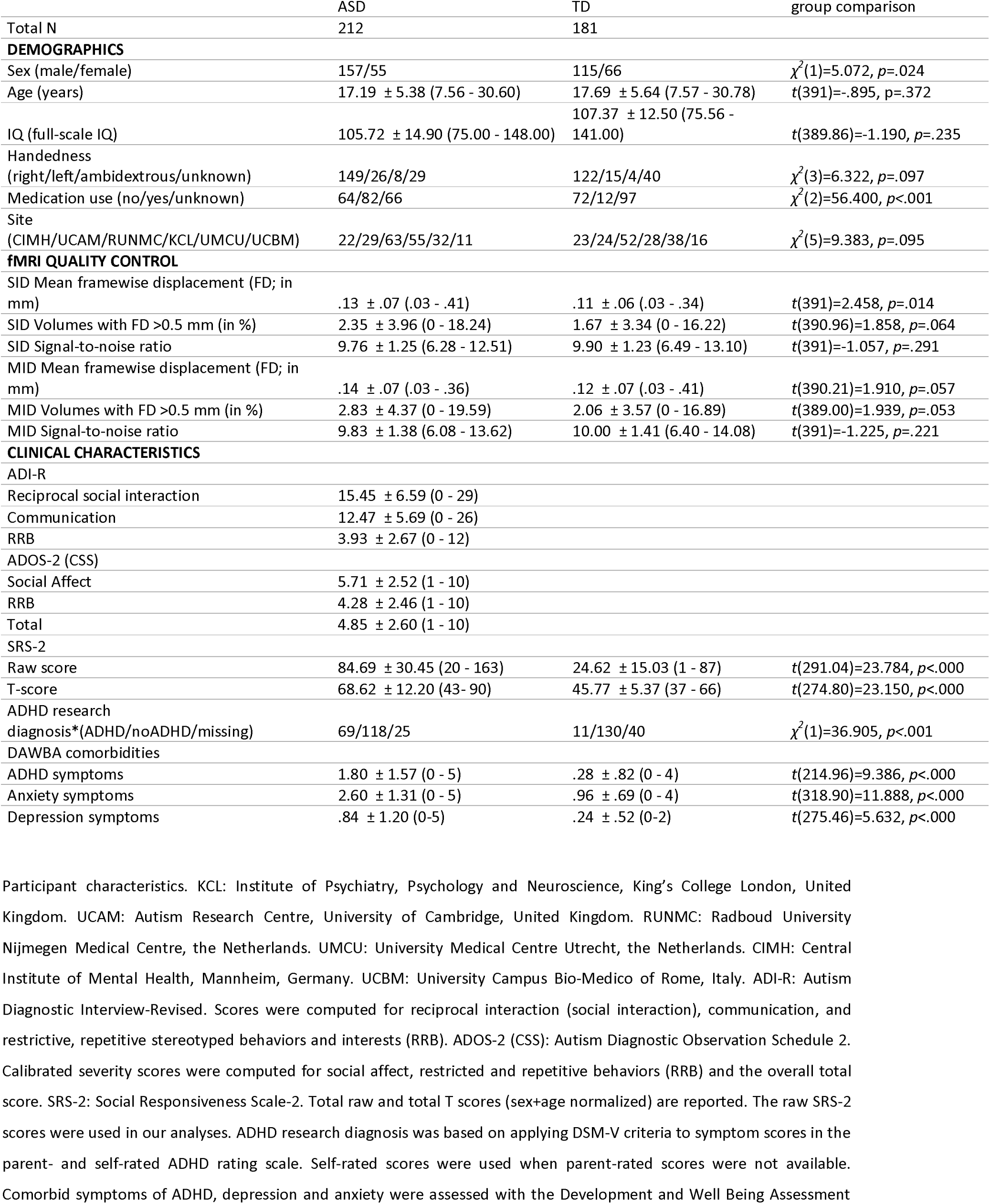
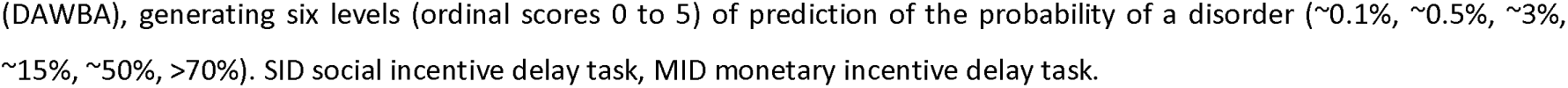
Sample characteristics.

|  | ASD | TD | group comparison |
| --- | --- | --- | --- |
| Total N | 212 | 181 |  |
| **DEMOGRAPHICS** |  |  |  |
| Sex (male/female) | 157/55 | 115/66 | $\chi^2(1)=5.072, p=.024$ |
| Age (years) | 17.19 ± 5.38 (7.56 - 30.60) | 17.69 ± 5.64 (7.57 - 30.78) | $t(391)=-.895, p=.372$ |
| IQ (full-scale IQ) | 105.72 ± 14.90 (75.00 - 148.00) | 107.37 ± 12.50 (75.56 - 141.00) | $t(389.86)=-1.190, p=.235$ |
| Handedness (right/left/ambidextrous/unknown) | 149/26/8/29 | 122/15/4/40 | $\chi^2(3)=6.322, p=.097$ |
| Medication use (no/yes/unknown) | 64/82/66 | 72/12/97 | $\chi^2(2)=56.400, p<.001$ |
| Site (CIMH/UCAM/RUNMC/KCL/UMCU/UCBM) | 22/29/63/55/32/11 | 23/24/52/28/38/16 | $\chi^2(5)=9.383, p=.095$ |
| **fMRI QUALITY CONTROL** |  |  |  |
| SID Mean framewise displacement (FD; in mm) | .13 ± .07 (.03 - .41) | .11 ± .06 (.03 - .34) | $t(391)=2.458, p=.014$ |
| SID Volumes with FD>0.5 mm (in %) | 2.35 ± 3.96 (0 - 18.24) | 1.67 ± 3.34 (0 - 16.22) | $t(390.96)=1.858, p=.064$ |
| SID Signal-to-noise ratio | 9.76 ± 1.25 (6.28 - 12.51) | 9.90 ± 1.23 (6.49 - 13.10) | $t(391)=-1.057, p=.291$ |
| MID Mean framewise displacement (FD; in mm) | .14 ± .07 (.03 - .36) | .12 ± .07 (.03 - .41) | $t(390.21)=1.910, p=.057$ |
| MID Volumes with FD>0.5 mm (in %) | 2.83 ± 4.37 (0 - 19.59) | 2.06 ± 3.57 (0 - 16.89) | $t(389.00)=1.939, p=.053$ |
| MID Signal-to-noise ratio | 9.83 ± 1.38 (6.08 - 13.62) | 10.00 ± 1.41 (6.40 - 14.08) | $t(391)=-1.225, p=.221$ |
| **CLINICAL CHARACTERISTICS** |  |  |  |
| ADI-R |  |  |  |
| Reciprocal social interaction | 15.45 ± 6.59 (0 - 29) |  |  |
| Communication | 12.47 ± 5.69 (0 - 26) |  |  |
| RRB | 3.93 ± 2.67 (0 - 12) |  |  |
| ADOS-2 (CSS) |  |  |  |
| Social Affect | 5.71 ± 2.52 (1 - 10) |  |  |
| RRB | 4.28 ± 2.46 (1 - 10) |  |  |
| Total | 4.85 ± 2.60 (1 - 10) |  |  |
| SRS-2 |  |  |  |
| Raw score | 84.69 ± 30.45 (20 - 163) | 24.62 ± 15.03 (1 - 87) | $t(291.04)=23.784, p<.000$ |
| T-score | 68.62 ± 12.20 (43 - 90) | 45.77 ± 5.37 (37 - 66) | $t(274.80)=23.150, p<.000$ |
| ADHD research diagnosis*(ADHD/noADHD/missing) | 69/118/25 | 11/130/40 | $\chi^2(1)=36.905, p<.001$ |
| DAWBA comorbidities |  |  |  |
| ADHD symptoms | 1.80 ± 1.57 (0 - 5) | .28 ± .82 (0 - 4) | $t(214.96)=9.386, p<.000$ |
| Anxiety symptoms | 2.60 ± 1.31 (0 - 5) | .96 ± .69 (0 - 4) | $t(318.90)=11.888, p<.000$ |
| Depression symptoms | .84 ± 1.20 (0-5) | .24 ± .52 (0-2) | $t(275.46)=5.632, p<.000$ |

Participant characteristics. KCL: Institute of Psychiatry, Psychology and Neuroscience, King's College London, United Kingdom. UCAM: Autism Research Centre, University of Cambridge, United Kingdom. RUNMC: Radboud University Nijmegen Medical Centre, the Netherlands. UMCU: University Medical Centre Utrecht, the Netherlands. CIMH: Central Institute of Mental Health, Mannheim, Germany. UCBM: University Campus Bio-Medico of Rome, Italy. ADI-R: Autism Diagnostic Interview-Revised. Scores were computed for reciprocal interaction (social interaction), communication, and restrictive, repetitive stereotyped behaviors and interests (RRB). ADOS-2 (CSS): Autism Diagnostic Observation Schedule 2. Calibrated severity scores were computed for social affect, restricted and repetitive behaviors (RRB) and the overall total score. SRS-2: Social Responsiveness Scale-2. Total raw and total T scores (sex+age normalized) are reported. The raw SRS-2 scores were used in our analyses. ADHD research diagnosis was based on applying DSM-V criteria to symptom scores in the parent- and self-rated ADHD rating scale. Self-rated scores were used when parent-rated scores were not available. Comorbid symptoms of ADHD, depression and anxiety were assessed with the Development and Well Being Assessment
(DAWBA), generating six levels (ordinal scores 0 to 5) of prediction of the probability of a disorder (~0.1%, ~0.5%, ~3%, ~15%, ~50%, >70%). SID social incentive delay task, MID monetary incentive delay task.

#### Clinical measures

Participants in the ASD group had an existing clinical diagnosis of ASD according to the DSM-IV/ICD-10 or DSM-5 criteria. ASD symptoms were comprehensively assessed using the Autism Diagnostic Interview-Revised (ADI-R; (20)) and Autism Diagnostic Observation Schedule 2 (ADOS-2; (21)) within the ASD group. We used the total raw score on the Social Responsiveness Scale Second Edition (SRS-2; (22)) to assess continuous autism traits, which was available across the study sample. Parent-rated scores were collected for ASD and TD individuals except for TD adults where only the self-report was assessed. We used self-rated scores wherever parent-rated scores were not available. Parent- or self-report of a psychiatric disorder was an exclusion criterion for the TD group. Information on the presence of a confirmed diagnosis of ADHD was not available in our sample. As a proxy, we estimated diagnostic status by applying DSM-5 criteria based on symptom scores collected with the parent- and self-rated ADHD DSM-5 rating scales (23).

#### Experimental paradigm

We adapted a social and a monetary incentive delay task (SID, MID) (4) as part of a reliable task battery (18, 24, 25). For details see figure 1 and supplementary material. SID and MID were collected as separate paradigms and combined during data analysis. SID was always presented first, followed by MID. The fMRI scanning session was preceded by a training session outside the MRI to ensure that all participants understood the task.

**Figure 1:**
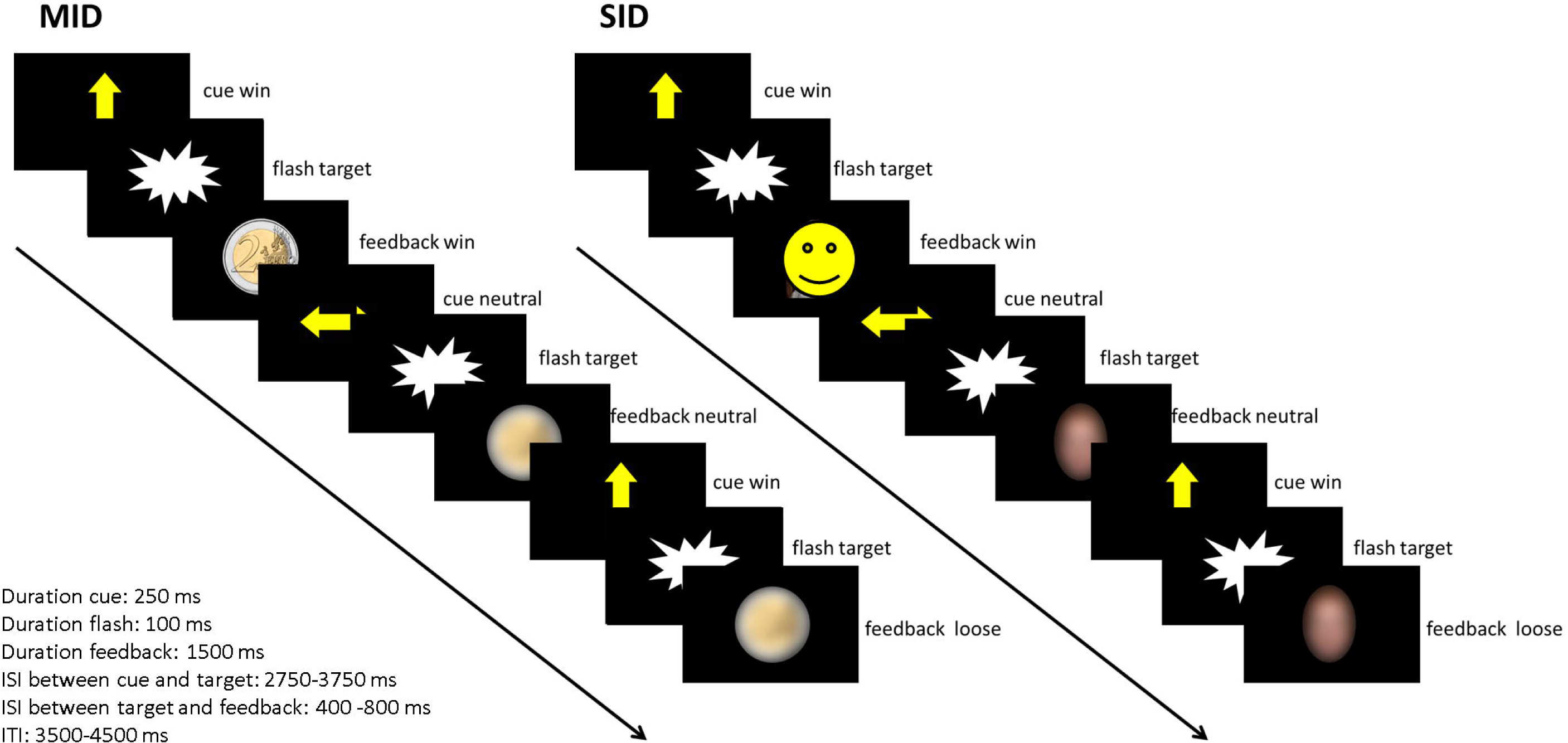
Task design of the monetary incentive delay task (MID) and social incentive delay task (SID). Participants were asked to give a speeded response (button press) to a visual target (screenflash). A cue arrow pointing upwards indicated the possibility to obtain a reward if responses were given within a predefined response time window (win trial). No reward option was given in trials preceded by a horizontal cue arrow (neutral trial). Sufficiently fast responses on win trials were followed by the presentation of a 2€/2£ coin in the MID and a smiling female face in the SID as feedback. Please note that due to BioRxiv policy, the actual face stimulus had to be replaced by a smiley in figure 1. Blurred control stimuli were presented in neutral trials and as feedback following slow responses in win trials. Cue presentation represents reward anticipation phase, while feedback presentation represents reward delivery phase. Note that the feedback presentation was temporally decoupled from the target presentation but not from the button press.

#### fMRI data acquisition

Functional MRI data were acquired on 3T scanners from different manufacturers (General Electric, Philips, Siemens) and harmonized as much as possible across sites (for details see supplementary material). Functional images were acquired using a BOLD-sensitive T2*-weighted echo-planar imaging (EPI) sequence (repetition time (TR) = 2 s, echo time (TE) = 30 ms, flip angle = 80°, matrix: 64 × 64, FOV: 192 × 192 mm, in-plane resolution: 3 x 3 mm, slice thickness: 4 mm, gap: 1 mm, 28 axial slices). A total of 151 volumes were obtained for each task, oriented approximately 20° steeper than the AC-PC-plane.

## Data analysis

### fMRI data preprocessing

Image preprocessing followed standard processing routines in SPM12 (http://www.fil.ion.ucl.ac.uk/spm/), including a two-pass realignment procedure, slice time correction, registration of the functional mean image to the Montreal Neurological Institute (MNI) template and spatial normalization into standard stereotactic space, application of resulting normalization parameters to the functional time series, resampling to 3 mm isotropic voxels, and smoothing with an 8 mm full-width at half-maximum Gaussian Kernel.

### Whole-brain level fMRI data analysis

SID and MID tasks were combined as two sessions in a general linear model (GLM) on the single subject level (see supplementary material for details). Within-subject effects were addressed at the subject-level by quantifying within subject effects of condition as differential response to win cues as compared to neutral cues for reward anticipation and differential response to successful win compared to neutral trials for reward delivery. Additionally, to quantify differential reward-specific responses between tasks, a contrast image for the interaction between condition (win, neutral) and task (SID, MID) was calculated.

Based on within-subject contrasts we assessed reward-specific brain activation (within subject effect of condition) and differential reward-specific responses between tasks (within-subject interaction condition x task) across the entire sample and tested for between-group differences. Contrast images were subjected to second-level GLMs with between-subject factor group (ASD vs. TD) and covariates age, sex, and site. The impact of ADHD comorbidity was explored in a separate model, where the ASD group was split into subgroups with (n=69) and without (n=118) comorbid ADHD based on estimated diagnostic status (ASD_+ADHD_ and ASD_-ADHD_, respectively) and compared to TD. TD individuals with elevated ADHD scores were excluded from this analysis. To assess the effect of autism traits, SRS-2 raw scores were added as additional covariate of interest in a separate model. Note that diagnostic group was accounted for in this model, ensuring that effects were not driven by differences in group means. To explore group differences on a whole-brain level, significance was defined as p_FWE_ < .05 with a cluster threshold of k≥5, peak-level corrected across the whole brain.

### Region of interest analysis

To increase sensitivity for putative case-control differences in the VS, we performed region of interest (ROI) analysis within a well-established a-priori defined bilateral mask of the VS (18). Mean contrast estimates (CE, contrasts cue win>cue neutral and successful win>neutral) for each participant and both tasks were extracted and analyzed using SPSS Software package (Version 25, IBM Corp., Armonk, NY, USA). Separate repeated measures ANOVAs with the within-subject factor task (MID, SID) and between-subject factor diagnosis (TD, ASD), and covariates age (mean-centered), sex, and study sites (dummy coded), were used to assess group differences for both reward processing phases (anticipation, delivery) in the left and right VS. To correct for investigating left and right VS activity separately, the critical alpha threshold was adjusted to p<.025 based on the Bonferroni procedure. Additionally, Bonferroni-correction was applied to all post-hoc pairwise comparisons. To assess the effect of autism traits, SRS-2 raw scores were added as additional covariate of interest in a separate model. Interaction terms between diagnosis and SRS-2 were added as well. The impact of ADHD comorbidity was explored in another separate model, where the between-subject factor diagnosis comprised three levels (TD, ASD_+ADHD_ and ASD_-ADHD_).

## Results

### Functional activation analysis

#### Reward anticipation

##### Whole-brain level analysis

Reward-specific brain activation was observed in an extensive network with peak activations in the bilateral VS, ACC/SMA, Thalamus, bilateral precentral gyrus and bilateral anterior insula/IFG for the anticipation of win compared to neutral trials collapsed across both reward tasks.

Reward-specific brain activation differed between diagnostic groups at the whole-brain level in the right VS (*F*_(1,384)_=22.84, *p*_FWE_=.017, *k*=8) during reward anticipation. A post-hoc T-test showed that activation was reduced in ASD compared to TD individuals.

See figure 2 A and B and table 2 for details.

**Table 2:**
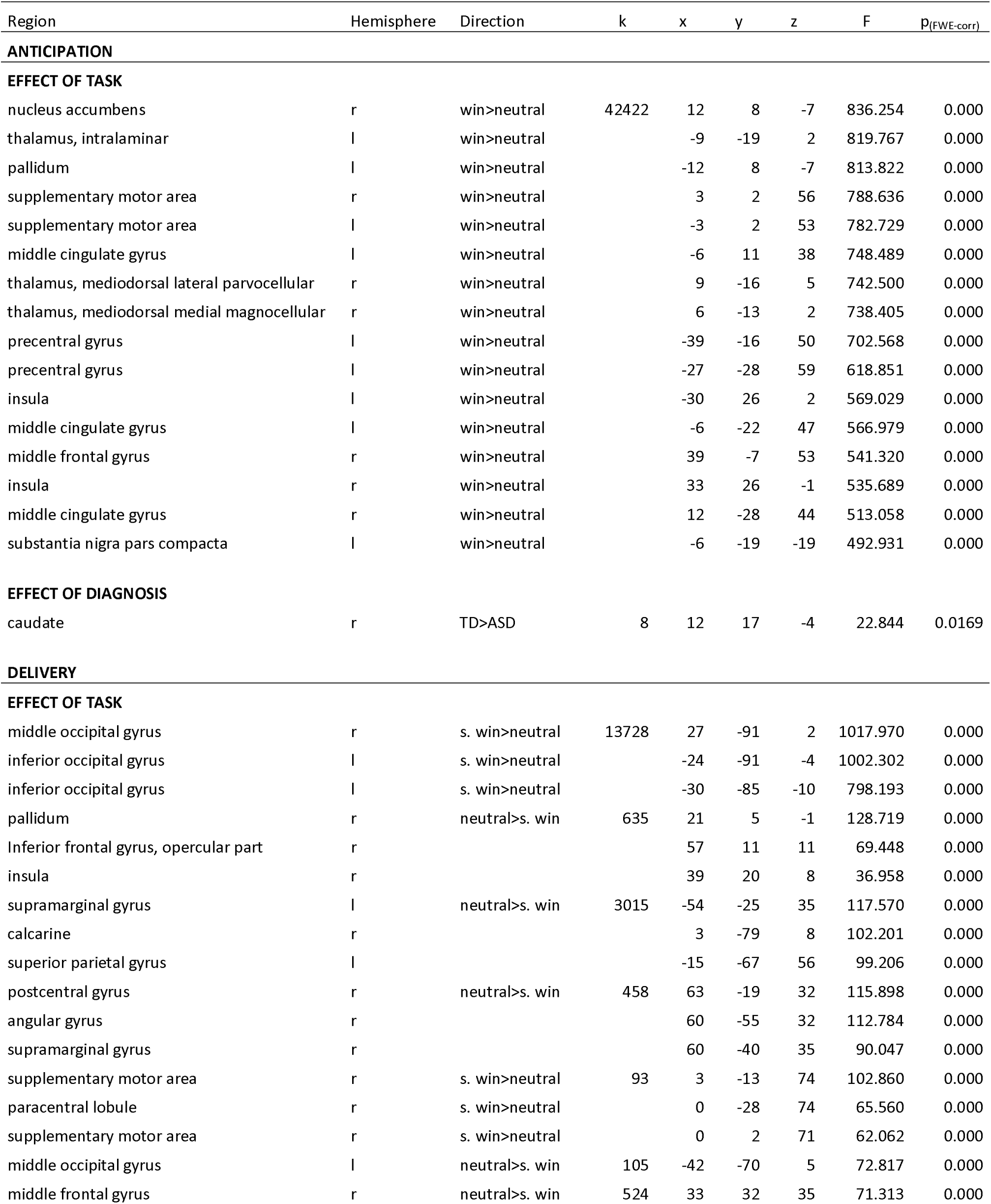
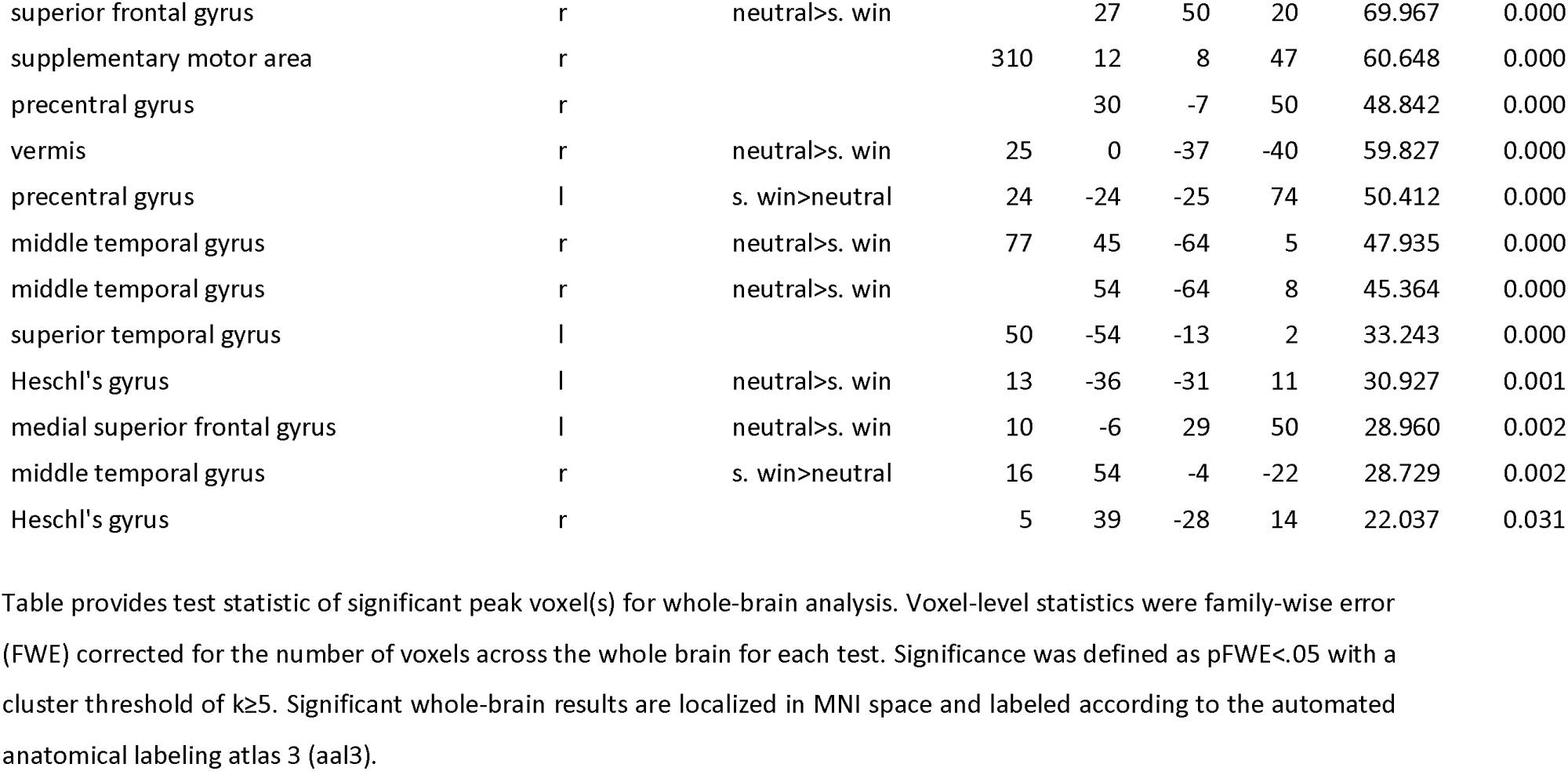
Whole-brain effects of brain activation during reward anticipation and delivery.

**Figure 2:**
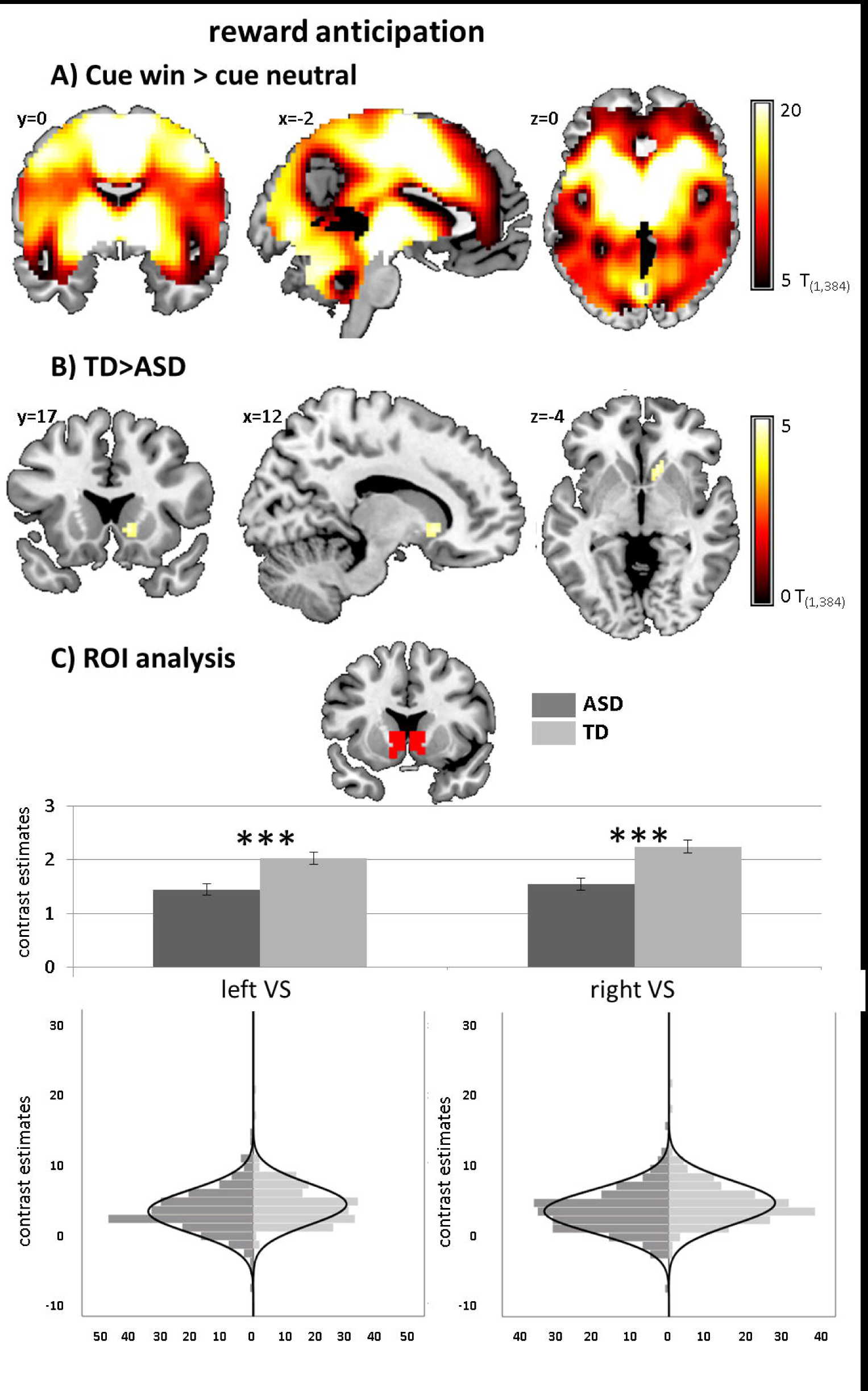
Brain activation to win compared to neutral cues. A) Whole-brain familywise error corrected activation across both ASD and TD individuals. B) Whole-brain familywise error corrected effect of diagnosis in the right ventral striatum C) Effect of diagnosis in the region of interest (ROI) analysis of the left and right ventral striatum with corresponding distribution plots. \*\*\**p*<.001. Distributions of ROI activation in cases and controls were compared using the Kolmogorov-Smirnov test, which suggested unequal distributions (left VS: *D*_(212,181)_=.156, *p*=.017; right VS: *D*_(212,181)_=.193, *p*=.001).

Differential reward-specific responses between tasks yielded activation in a network with peak activations in the bilateral VS, anterior cingulate cortex (ACC)/supplementary motor area (SMA), thalamus, bilateral precentral gyrus and bilateral anterior insula/inferior frontal gyrus (IFG) (see fig 3A and table 3). Post-hoc T-tests showed stronger differential activation in the MID compared to the SID across all these regions.

**Table 3:** Whole-brain effects of brain activation for interaction between cue (win, neutral) and task (SID, MID).

| Region | Hemisphere | Direction | k | x | y | z | F | $p_{(FWE-corr)}$ |
| --- | --- | --- | --- | --- | --- | --- | --- | --- |
| **ANTICIPATION** |  |  |  |  |  |  |  |  |
| **INTERACTION TASK\*CUE** |  |  |  |  |  |  |  |  |
| supplementary motor area | r | MID>SID | 29325 | 3 | -4 | 62 | 157.583 | 0.000 |
| supplementary motor area | l | MID>SID |  | -3 | -7 | 62 | 157.215 | 0.000 |
| supplementary motor area | r | MID>SID |  | 6 | 2 | 56 | 153.182 | 0.000 |
| precentral gyrus | l | MID>SID |  | -36 | -13 | 53 | 138.596 | 0.000 |
| caudate | r | MID>SID |  | 9 | 8 | -1 | 132.749 | 0.000 |
| supplementary motor area | l | MID>SID |  | -6 | 2 | 50 | 129.527 | 0.000 |
| pallidum | l | MID>SID |  | -12 | 5 | 2 | 127.187 | 0.000 |
| precentral gyrus | l | MID>SID |  | -30 | -28 | 59 | 119.560 | 0.000 |
| precuneus | l | MID>SID |  | -12 | -73 | 47 | 118.639 | 0.000 |
| precentral gyrus | l | MID>SID |  | -24 | -25 | 59 | 118.186 | 0.000 |
| postcentral gyrus | l | MID>SID |  | -30 | -43 | 65 | 118.030 | 0.000 |
| superior parietal gyrus | l | MID>SID |  | -27 | -49 | 65 | 117.861 | 0.000 |
| precentral gyrus | l | MID>SID |  | -21 | -19 | 65 | 115.833 | 0.000 |
| precentral gyrus | l | MID>SID |  | -27 | -10 | 65 | 109.229 | 0.000 |
| thalamus, mediodorsal lateral parvocellular | l | MID>SID |  | -9 | -16 | 5 | 107.077 | 0.000 |
| thalamus, ventral lateral | l | MID>SID |  | -12 | -13 | 2 | 105.532 | 0.000 |
| inferior temporal gyrus | l | MID>SID | 7 | -42 | -28 | -19 | 24.929 | 0.009 |
| **DELIVERY** |  |  |  |  |  |  |  |  |
| **INTERACTION TASK\*CUE** |  |  |  |  |  |  |  |  |
| pallidum | l | MID>SID | 20268 | -18 | 5 | -4 | 923.157 | 0.000 |
| thalamus, intralaminar | l | MID>SID |  | -6 | -19 | -4 | 905.845 | 0.000 |
| supplementary motor area | l | MID>SID |  | -6 | -1 | 56 | 862.060 | 0.000 |
| Calcarine fissure and surrounding cortex | r | MID>SID | 426 | 18 | -70 | 11 | 120.088 | 0.000 |
| cuneus | r | MID>SID |  | 18 | -70 | 38 | 51.291 | 0.000 |
| Calcarine fissure and surrounding cortex | l | MID>SID | 396 | -15 | -73 | 11 | 108.793 | 0.000 |
| precuneus | l | MID>SID |  | -15 | -70 | 35 | 43.234 | 0.000 |
| insula | r | SID>MID | 217 | 45 | -10 | 11 | 96.633 | 0.000 |
| rolandic operculum | r | SID>MID |  | 63 | -7 | 8 | 72.488 | 0.000 |
| postcentral gyrus | r | SID>MID |  | 60 | -4 | 26 | 70.168 | 0.000 |
| postcentral gyrus | r | SID>MID | 489 | 48 | -34 | 62 | 89.013 | 0.000 |
| angular gyrus | r | SID>MID |  | 42 | -64 | 41 | 75.999 | 0.000 |
| superior parietal gyrus | r | SID>MID |  | 42 | -49 | 62 | 60.413 | 0.000 |
| inferior frontal gyrus, pars orbitalis | l | SID>MID | 45 | -48 | 32 | -10 | 81.258 | 0.000 |
| posterior orbital gyrus | l | SID>MID |  | -42 | 26 | -16 | 45.152 | 0.000 |
| inferior frontal gyrus, triangular part | l | SID>MID |  | -51 | 29 | -1 | 41.552 | 0.000 |
| posterior orbital gyrus | r |  | 43 | 27 | 11 | -25 | 51.957 | 0.000 |
| middle frontal gyrus | l | SID>MID | 47 | -42 | 11 | 53 | 42.081 | 0.000 |
| middle temporal gyrus | r | SID>MID | 23 | 60 | -10 | -22 | 41.604 | 0.000 |
| middle frontal gyrus | r | SID>MID | 34 | 39 | 56 | -4 | 41.152 | 0.000 |
| temporale pole, middle temporal gyrus | l |  | 27 | -39 | 5 | -19 | 33.684 | 0.000 |
| inferior parietal gyrus | r | MID>SID | 5 | 57 | -37 | 47 | 33.175 | 0.000 |
| postcentral gyrus | l | SID>MID | 12 | -48 | -40 | 59 | 32.283 | 0.000 |
| superior parietal gyrus | r | MID>SID | 18 | 30 | -49 | 47 | 32.169 | 0.000 |
| vermis | l | SID>MID | 7 | -3 | -46 | -28 | 28.712 | 0.002 |
| precuneus | l | MID>SID | 5 | -27 | -52 | 14 | 26.502 | 0.005 |
| lateral orbital gyrus | r | SID>MID | 5 | 45 | 38 | -16 | 26.407 | 0.005 |

Table provides test statistic of significant peak voxel(s) for whole-brain analysis. Voxel-level statistics were family-wise error (FWE) corrected for the number of voxels across the whole brain for each test. Significance was defined as $p_{FWE} < .05$ with a cluster threshold of $k \geq 5$ . Significant whole-brain results are localized in MNI space and labeled according to the automated anatomical labeling atlas 3 (aal3). Direction of the effect tested post-hoc via t-tests.

**Figure 3:**
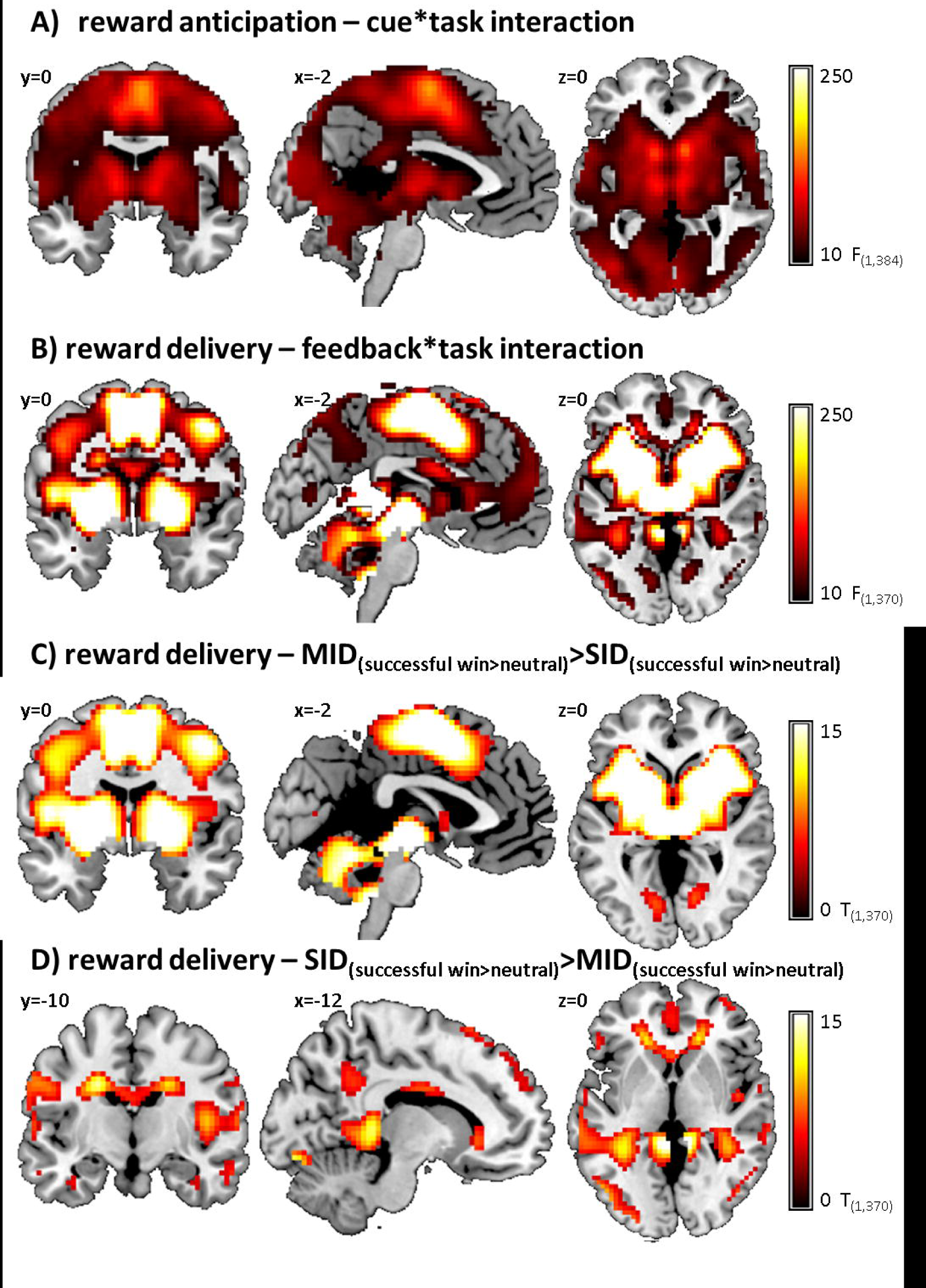
A) Interaction effect of cue (win, neutral) and task (SID, MID) indicating higher differential activation in MID compared to SID. B) Interaction effect of feedback (win, neutral) and task (SID, MID) C) Differentially increased activity in MID compared to SID for successful win compared to neutral trials. D) Differentially increased activity in SID compared to MID for successful win compared to neutral trials.

Differential reward-specific responses between tasks did not differ between individuals with and without ASD, however, we report differences between ASD and TD in the SID and MID separately in the supplementary material (figure S1 and tables S2 and S3) to allow for a comparison with previous studies.

#### ROI analysis

Individuals with ASD differed from TD individuals on average regarding reward-specific brain activation within the left (*F*_(1,384)_ =14.163, *p*<.001, partial *η*^*2*^=.036) and right (*F* _(1,384)_=18.693, *p*<.001, partial *η*^*2*^=.046) VS ROI with reduced activation in ASD (left: *M*=1.45, *SD*=1.53, right: *M*=1.54, *SD*=1.58) compared to TD individuals (left: *M*=2.03, *SD*=1.53, *d*=−0.39, right: *M*=2.25, *SD*=1.59, *d*=−0.44). See figure 3C. There was no significant interaction between diagnosis and task (left VS: *F*_(1,384)_ =2.754, *p*=.098, partial *η*^*2*^=.007, right VS: *F* _(1,384)_ =2.999, *p*=.084, partial *η*^*2*^=.008).

#### Reward delivery

##### Whole-brain level analysis

During reward delivery, collapsed across both reward tasks, the feedback of successful win compared to neutral trials activated a network with peak activations in the visual cortex, ACC/SMA, thalamus, bilateral precentral gyrus and bilateral anterior insula/IFG, while reduced activation in comparison to neutral trials was observed in a network comprising occipital, frontal and temporal regions as well as the thalamus and the bilateral pallidum.

There was no significant effect of diagnostic group on reward-specific brain activation at the whole-brain level. See figure 4A and table 2 for details.

**Figure 4:**
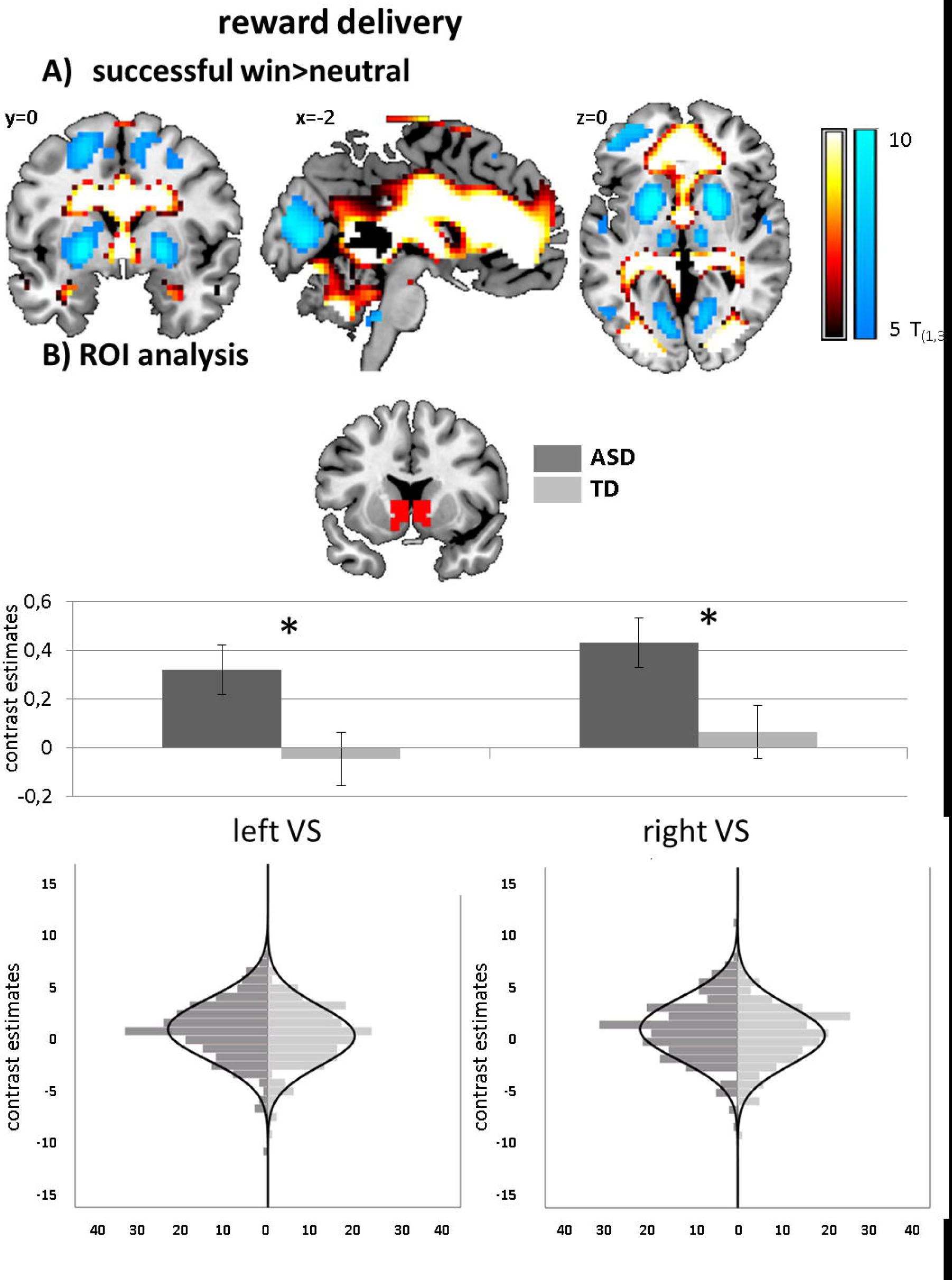
Brain activation to reward delivery. A) Whole-brain familywise error corrected activation increase (warm colours) and decrease (cold colours) to successful win compared to neutral trials across both ASD and TD individuals. B) Effect of diagnosis in the region of interest (ROI) analysis of the left and right ventral striatum with corresponding distribution plots. \**p*<.05. Distributions of ROI activation in cases and controls were compared using the Kolmogorov-Smirnov test, which suggested no evidence for unequal distributions (left VS: *D*_(205,174)_=.120, *p*=.134; right VS: *D*_(205,174)_=.112, *p*=.190).

Differential reward-specific responses between tasks showed activation in a network with peak activations in the bilateral VS, ACC/SMA, thalamus, left precentral gyrus and bilateral anterior insula/IFG (see fig 3B and table 3). Subsequent T-tests indicated stronger differential activation in the MID compared to the SID in these peak regions (see figure 3C), while stronger differential activation in the SID compared to the MID was found in a network with peak activations in bilateral hippocampus, bilateral fusiform gyrus, bilateral lingual gyrus and ACC (see figure 3D).

The interaction effect of diagnosis was not significant for differential reward-specific responses between tasks. However, we report differences between ASD and TD in the SID and MID separately in the supplementary material (figure S2 and tables S2 and S3) to allow comparison with previous studies.

##### ROI analysis

The difference regarding reward-specific brain activation between ASD and TD individuals within the left (*F*_(1,370)_ =4.829, *p*=.029, partial *η*^*2*^=.013) and right VS (*F* _(1,370)_ =4.719, *p*=.030, partial *η*^*2*^=.013) yielded increased activation in ASD (left: *M*=.31, *SD*=1.46, right: *M*=.42, *SD*=1.47) compared to TD (left: *M*=−.02, *SD*=1.46, *d*=0.23, right: *M*=.09, *SD*=1.47, *d*=0.23) but did not survive correction for multiple comparisons. See figure 4C. There was no significant interaction between diagnosis and task (left VS: *F*_(1,370)_ =1.057, *p*=.304, partial *η*^*2*^=.003, right VS: *F* _(1,370)_ =1.684, *p*=.195, partial *η*^*2*^=.005).

#### Dimensional effects

For both reward anticipation and delivery there was no significant main effect of autism trait scores and no interaction between diagnosis and autism trait scores in the VS or on the whole-brain level. Statistics are summarized in table 4. Autism trait scores also showed no significant effect when analyzing TD and ASD individuals separately.

**Table 4:**
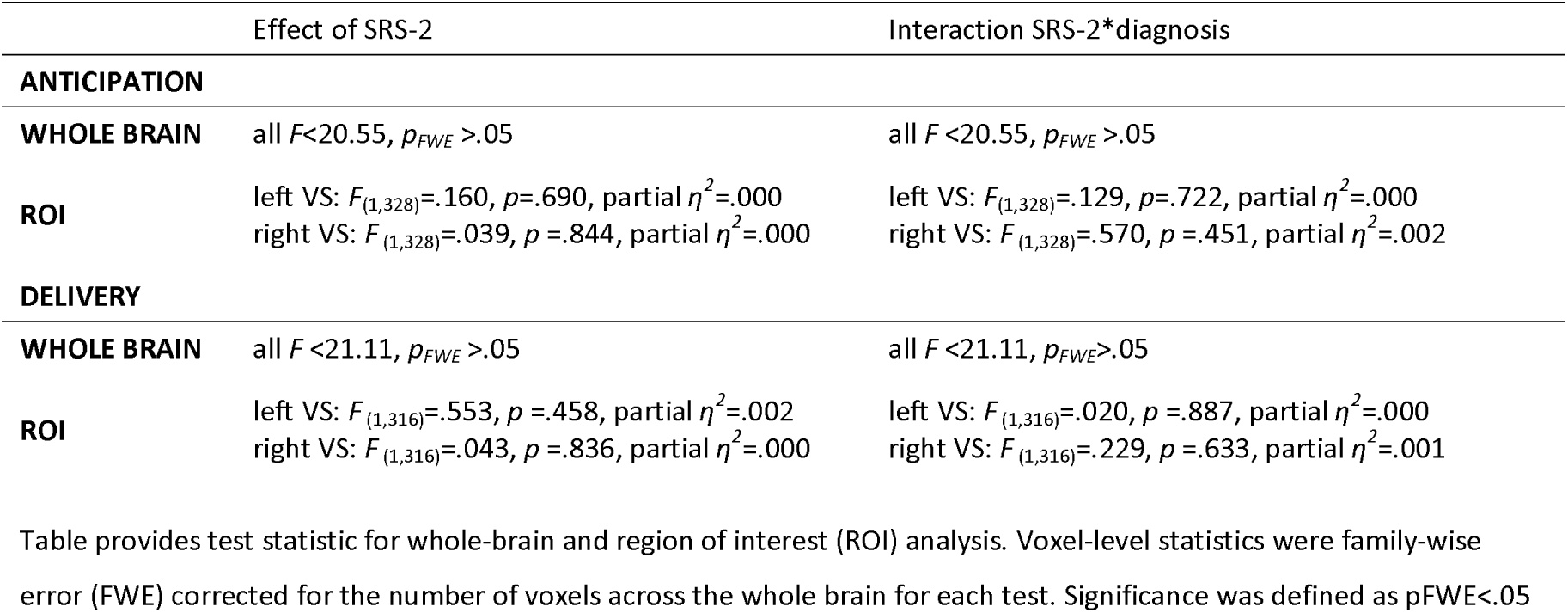
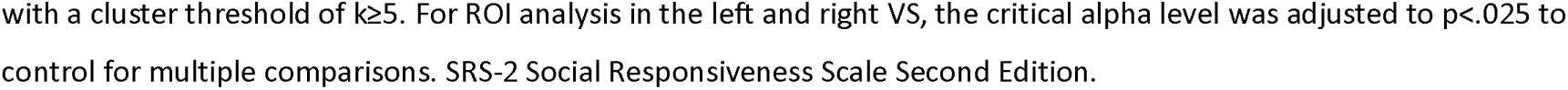
Whole-brain and region of interest (ROI) effects of autism trait-related brain activation during reward anticipation and delivery.

|  | Effect of SRS-2 | Interaction SRS-2*diagnosis |
| --- | --- | --- |
| **ANTICIPATION** |  |  |
| **WHOLE BRAIN** | all $F < 20.55$, $p_{FWE} > .05$ | all $F < 20.55$, $p_{FWE} > .05$ |
| **ROI** | left VS: $F_{(1,328)} = .160$, $p = .690$, partial $\eta^2 = .000$<br>right VS: $F_{(1,328)} = .039$, $p = .844$, partial $\eta^2 = .000$ | left VS: $F_{(1,328)} = .129$, $p = .722$, partial $\eta^2 = .000$<br>right VS: $F_{(1,328)} = .570$, $p = .451$, partial $\eta^2 = .002$ |
| **DELIVERY** |  |  |
| **WHOLE BRAIN** | all $F < 21.11$, $p_{FWE} > .05$ | all $F < 21.11$, $p_{FWE} > .05$ |
| **ROI** | left VS: $F_{(1,316)} = .553$, $p = .458$, partial $\eta^2 = .002$<br>right VS: $F_{(1,316)} = .043$, $p = .836$, partial $\eta^2 = .000$ | left VS: $F_{(1,316)} = .020$, $p = .887$, partial $\eta^2 = .000$<br>right VS: $F_{(1,316)} = .229$, $p = .633$, partial $\eta^2 = .001$ |

Table provides test statistic for whole-brain and region of interest (ROI) analysis. Voxel-level statistics were family-wise error (FWE) corrected for the number of voxels across the whole brain for each test. Significance was defined as $p_{FWE} < .05$
with a cluster threshold of $k \geq 5$ . For ROI analysis in the left and right VS, the critical alpha level was adjusted to $p < .025$ to control for multiple comparisons. SRS-2 Social Responsiveness Scale Second Edition.

#### Effect of ADHD comorbidity

During reward anticipation, ROI analysis yielded a significant effect of group in the left (*F*_(1,307)_=5.172, *p*=.006, partial *η*^*2*^=.032) and right (*F*_(1,307)_ =6.761, *p=*.001, partial *η*^*2*^=.042) VS (see figure 5 A). Pairwise comparisons revealed that this effect was driven by significantly increased VS activity in TD compared to ASD_-ADHD_ (left: *p*=.006, *d*=0.40, right: *p*=.001, *d*=0.46), while there was no significant difference between TD and ASD_+ADHD_ (left: *p*=.144, *d*=0.30, right: *p*=.099, *d*=0.32) or between the two ASD subgroups (left: *p*=1.000, *d*=−0.09, right: *p*=1.000, *d*=−0.13). For reward delivery, a borderline significant effect of group emerged in the right VS (*F* _(1,297)_=3.715, *p=*.026, partial *η*^*2*^=.024, see figure 5 B) with significantly increased VS activity in ASD_-ADHD_ compared to TD (*p*=.020, *d*=0.35) and no difference between TD and ASD_+ADHD_ (*p*=.741, *d*=−0.18) or between the two ASD subgroups (*p*=.810, *d*=0.17). Across both reward processing stages, there was no significant effect of group on the whole-brain level and no significant interaction with the type of reward (social, monetary).

**Figure 5:**
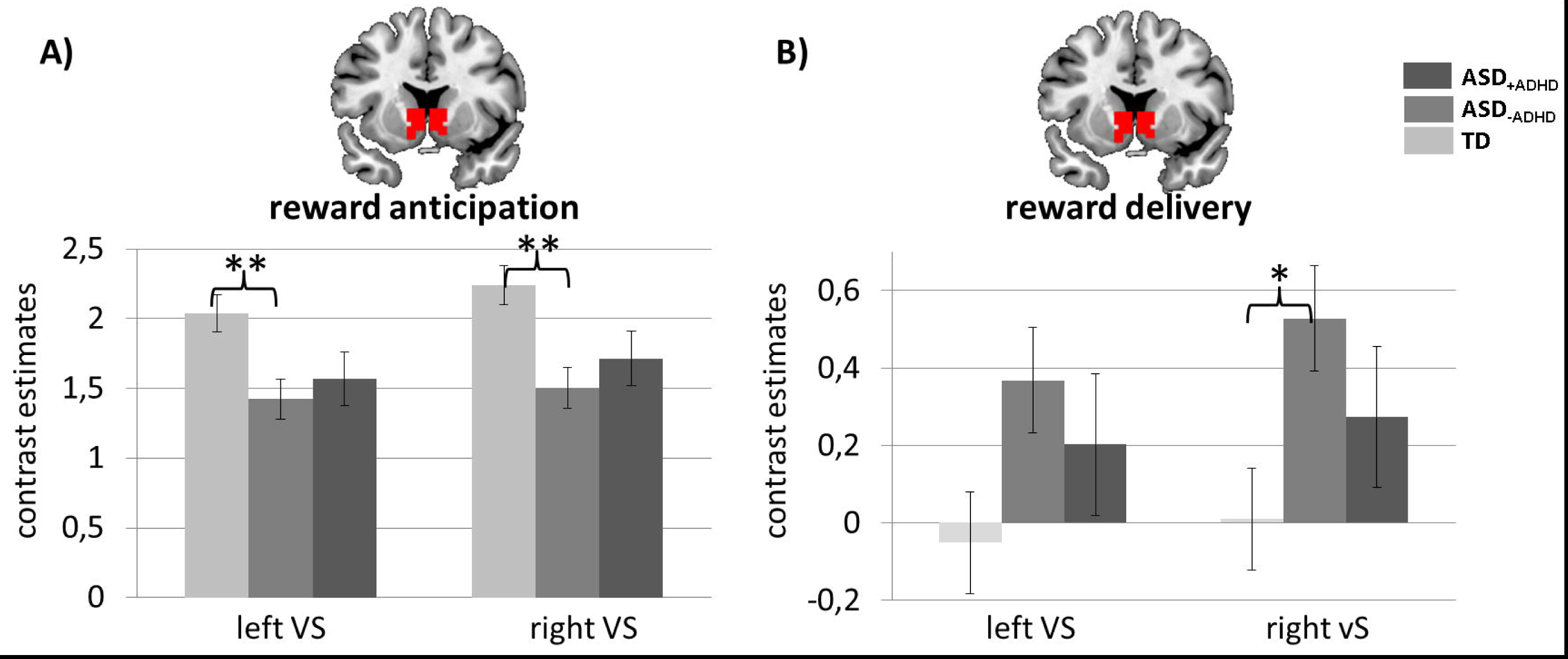
Contrast estimates for ventral striatal activation in individuals with ASD and elevated ADHD symptoms (ASD_+ADHD)_, individuals with ASD without elevated ADHD symptoms (ASD_-ADHD)_ and typically developing individuals without elevated ADHD symptoms (TD) during A) reward anticipation and B) delivery. \*\**p*<.01, \**p*<.05.

#### Control analyses

Supplemental control analyses showed that results were not systematically explained by head motion, acquisition site, handedness, sex, intelligence quotient (IQ) or medication status. Effects of age (linear and quadratic) were observed during reward delivery in the right superior medial frontal gyrus and the left amygdala, pallidum and (at trend-level) the VS, respectively. These effects did not differ between ASD and TD. For reward delivery, we were not able to replicate the effect of diagnosis when investigating only female participants, only right-handed participants, or when excluding participants from RUNMC or KCL. While this likely reflects decreased statistical power due to reduced subsample sizes, it also warrants further exploration of potential sources of heterogeneity in future studies. Details on the control analyses are provided in the supplementary material.

## Discussion

In the present study, we assessed functional brain activation during monetary and social reward anticipation and delivery in a well-powered sample comprising ASD and TD individuals. This allowed us to examine effects of reward type during both reward processing phases. We found a reduction of VS activity during reward anticipation in individuals with ASD that did not differ between social and monetary rewards. In contrast, during reward delivery, we found that increased VS activity in ASD compared to TD across both social and monetary reward conditions did not survive correction for multiple comparisons. These results do not support opposing effect of social and monetary reward types, but rather point towards a general hypoactivity of VS in ASD during anticipation of rewards. This is in contrast to the hypothesis of a predominantly social motivation deficit (1) and previous findings in a recent meta-analysis (13). We conclude that, in ASD, general hypoactivation during the anticipation of rewards indicates attenuated subjective reward value independent of social content. Our finding is in line with a previous study investigating negative social and monetary reinforcement (26) and extends beyond ASD to other conditions such as schizophrenia and bipolar disorder (27), pointing towards a potential shared motivational shift in these conditions that need further investigation.

Our results on reward delivery do not show substantial differences between ASD and TD individuals. While this is in contrast to meta-analytic findings (13) and previous studies showing striatal hypoactivity during monetary reward delivery (7-9), it is in line with studies showing no significant group differences during social reward (5, 6), monetary reward (5), or showing differences only at an uncorrected threshold (10). In summary, our results suggest that both monetary and social rewards are eliciting reward-related brain activity upon delivery that is not strikingly different in ASD and TD. Behaviorally, individuals with ASD did not differ from TD rewarding reaction times and accuracy (see supplementary material), which is in line with previous findings (6, 7, 10, 28-30).

The failure to sufficiently activate the reward system during anticipation of these rewards might suggest a disrupted feedback loop between “liking” a reward and “wanting” a subsequent reward in ASD compared to TD. This might suggest atypical reinforcement-dependent learning (31) and/or salience processing in ASD (32-35), irrespective of reward type, and does not support the idea of a reward processing deficit predominantly for social rewards, as proposed by the social motivation hypothesis (1). A hypothesis of generally atypical reinforcement-dependent learning in ASD is however challenged by studies reporting elevated reward system responsivity in ASD to stimuli of high interest for autistic individuals (6, 9). These findings indicate intact, possibly even hyperactive reinforcement-dependent learning when stimuli with high individual interest are involved. Future work is therefore needed, exploring potential changes in feedback loops underlying altered reinforcement-dependent learning in ASD using connectivity metrics (36-38) and different reward types, as well as exploring links to atypical salience processing in ASD (39-41).

While significant differences between diagnostic groups were found, we did not observe significant associations between autism trait scores (SRS-2) and functional brain activation across the whole sample or within ASD and TD separately. Clements et al. (13) found a large (r=−.72) but non-significant association between SRS scores and activity in the caudate, with decreased activity associated with increased symptom severity for social reward types only. In supplemental analyses (see supplementary table S2) we assessed effects of autism trait scores for MID and SID separately, but observed no significant effect in this separate analysis. Although others found associations between dimensional autism measures and reward-related brain activity (5, 10, 29), our results are in line with previous studies also finding no association with dimensional autism measures (7, 9). Previous studies argued that their null findings might be due to insufficient power and insufficient range of scores in the ASD group (7, 9), which can be ruled out by the present findings.

Elevated levels of ADHD symptoms did not have an additive effect on reward system dysfunction in ASD, in contrast to our hypothesis. During reward anticipation, VS activity was reduced only in those individuals with ASD that had subthreshold levels of ADHD (ASD_-ADHD_) compared to TD, while those individuals with ASD that had elevated ADHD levels (ASD_+ADHD_,) did not differ significantly from TD or ASD_-ADHD_. During reward delivery, differences between the three groups were not strong enough to reach statistical significance when correcting for multiple comparisons. However, the direction of the effect also suggested the largest deviation for the ASD_-ADHD_ group. These results support an alternative hypothesis of ADHD symptoms partly balancing out ASD-related motivational deficits. This would be in line with previous findings, where individuals with ASD showed generally low VS reactivity, and individuals with ADHD showed high VS reactivity to both social and monetary reward types (10). However, this study did not differentiate between reward anticipation and delivery. While during monetary reward anticipation, VS hypoactivation is discussed as a fairly consistent finding in adults and adolescents with ADHD (15, but see 42), findings are more inconsistent in children (11, 43). For monetary reward delivery, increased VS activity in ADHD has been reported ((42, 44-46) but see (29)). Importantly, information on social reward processing in ADHD is scarce. Information on the presence of a confirmed diagnosis of ADHD was not available in our sample, and a questionnaire-derived proxy was used instead. This might have significantly impacted our findings, as ADHD-like behaviors might have been misclassified. As a consequence, our finding requires further investigation using clinically confirmed information on ASD-ADHD co-occurrence.

While the present study provides important insights into group-level, on-average reward processing alterations in autism, a number of limitations need to be addressed. First, while our findings of differences in reward processing between ASD and TD were significant for the anticipatory phase, effect sizes were small. This likely reflects substantial between-subject heterogeneity partly attributable to the multicenter design of the study and to the intention of collecting a representative dataset but most importantly reflecting the heterogeneity of ASD. We aim to further explore this heterogeneity within the LEAP sample using classification and stratification approaches (47) in future studies. Second, the task design did not allow for a neat separation between feedback presentation and motor response (short inter-stimulus interval, no jitter). Thus, we cannot rule out that findings in the delivery phase were influenced by motor activity.

In summary, the present study demonstrates significant reduction of VS activity during the anticipation of rewards in individuals with ASD irrespective of the type of reward, and subthreshold hyperactivity of VS during the delivery of these rewards. In contrast to our hypothesis, altered reward processing was not exacerbated by elevated ADHD symptoms. This might suggest generally atypical reward processing in ASD that is partly balanced out by co-occurring ADHD. This provides important insights, specifically as the impact of co-occurring ADHD has not been consistently assessed in previous studies on reward processing alterations in ASD and might contribute towards the heterogeneity of findings. Although further exploration of the underlying mechanisms is needed, the present study advances our understanding of the neuronal underpinnings of ASD by suggesting attenuated subjective reward value independent of social content and ADHD symptoms.

## Acknowledgements

This work was supported by EU-AIMS (European Autism Interventions), which received support from the Innovative Medicines Initiative Joint Undertaking under grant agreement no. 115300, the resources of which are composed of financial contributions from the European Union’ s Seventh Framework Programme (grant FP7/2007-2013), from the European Federation of Pharmaceutical Industries and Associations companies ‘in-kind contributions, and from Autism Speaks as well as AIMS-2-TRIALS which received support from the Innovative Medicines Initiative 2 Joint Undertaking under grant agreement No 777394. This joint undertaking receives support from the European Union’s Horizon 2020 research and innovation programme and EFPIA and AUTISM SPEAKS, Autistica, SFARI.

## References

1. Chevallier C, Kohls G, Troiani V, Brodkin ES, Schultz RT (2012): The social motivation theory of autism. Trends in cognitive sciences. 16:231–239.

2. Scott-Van Zeeland AA, Dapretto M, Ghahremani DG, Poldrack RA, Bookheimer SY (2010): Reward processing in autism. Autism research : official journal of the International Society for Autism Research. 3:53–67.

3. Haber SN, Knutson B (2010): The reward circuit: linking primate anatomy and human imaging. Neuropsychopharmacology : official publication of the American College of Neuropsychopharmacology. 35:4–26.

4. Delmonte S, Balsters JH, McGrath J, Fitzgerald J, Brennan S, Fagan AJ, et al. (2012): Social and monetary reward processing in autism spectrum disorders. Molecular autism. 3:7.

5. Dichter GS, Richey JA, Rittenberg AM, Sabatino A, Bodfish JW (2012): Reward circuitry function in autism during face anticipation and outcomes. Journal of autism and developmental disorders. 42:147–160.

6. Kohls G, Antezana L, Mosner MG, Schultz RT, Yerys BE (2018): Altered reward system reactivity for personalized circumscribed interests in autism. Molecular autism. 9:9.

7. Kohls G, Schulte-Ruther M, Nehrkorn B, Muller K, Fink GR, Kamp-Becker I, et al. (2013): Reward system dysfunction in autism spectrum disorders. Social cognitive and affective neuroscience. 8:565–572.

8. Assaf M, Hyatt CJ, Wong CG, Johnson MR, Schultz RT, Hendler T, et al. (2013): Mentalizing and motivation neural function during social interactions in autism spectrum disorders. NeuroImage Clinical. 3:321–331.

9. Dichter GS, Felder JN, Green SR, Rittenberg AM, Sasson NJ, Bodfish JW (2012): Reward circuitry function in autism spectrum disorders. Social cognitive and affective neuroscience. 7:160–172.

10. Kohls G, Thonessen H, Bartley GK, Grossheinrich N, Fink GR, Herpertz-Dahlmann B, et al. (2014): Differentiating neural reward responsiveness in autism versus ADHD. Developmental cognitive neuroscience. 10:104–116.

11. van Hulst BM, de Zeeuw P, Bos DJ, Rijks Y, Neggers SF, Durston S (2017): Children with ADHD symptoms show decreased activity in ventral striatum during the anticipation of reward, irrespective of ADHD diagnosis. Journal of child psychology and psychiatry, and allied disciplines. 58:206–214.

12. Lai MC, Lombardo MV, Chakrabarti B, Baron-Cohen S (2013): Subgrouping the autism “spectrum”: reflections on DSM-5. PLoS Biol. 11:e1001544.

13. Clements CC, Zoltowski AR, Yankowitz LD, Yerys BE, Schultz RT, Herrington JD (2018): Evaluation of the Social Motivation Hypothesis of Autism: A Systematic Review and Meta-analysis. JAMA psychiatry. 75:797–808.

14. Solberg BS, Zayats T, Posserud MB, Halmoy A, Engeland A, Haavik J, et al. (2019): Patterns of Psychiatric Comorbidity and Genetic Correlations Provide New Insights Into Differences Between Attention-Deficit/Hyperactivity Disorder and Autism Spectrum Disorder. Biological psychiatry. 86:587–598.

15. Plichta MM, Scheres A (2014): Ventral-striatal responsiveness during reward anticipation in ADHD and its relation to trait impulsivity in the healthy population: a meta-analytic review of the fMRI literature. Neurosci Biobehav Rev. 38:125–134.

16. Chantiluke K, Christakou A, Murphy CM, Giampietro V, Daly EM, Ecker C, et al. (2014): Disorder-specific functional abnormalities during temporal discounting in youth with Attention Deficit Hyperactivity Disorder (ADHD), Autism and comorbid ADHD and Autism. Psychiatry research. 223:113–120.

17. Loth E, Charman T, Mason L, Tillmann J, Jones EJH, Wooldridge C, et al. (2017): The EU-AIMS Longitudinal European Autism Project (LEAP): design and methodologies to identify and validate stratification biomarkers for autism spectrum disorders. Molecular autism. 8:24.

18. Plichta MM, Schwarz AJ, Grimm O, Morgen K, Mier D, Haddad L, et al. (2012): Test-retest reliability of evoked BOLD signals from a cognitive-emotive fMRI test battery. NeuroImage. 60:1746–1758.

19. Charman T, Loth E, Tillmann J, Crawley D, Wooldridge C, Goyard D, et al. (2017): The EU-AIMS Longitudinal European Autism Project (LEAP): clinical characterisation. Molecular autism. 8:27.

20. Lord C, Rutter M, Le Couteur A (1994): Autism Diagnostic Interview-Revised: a revised version of a diagnostic interview for caregivers of individuals with possible pervasive developmental disorders. Journal of autism and developmental disorders. 24:659–685.

21. Lord C, Risi S, Lambrecht L, Cook EH, Jr., Leventhal BL, DiLavore PC, et al. (2000): The autism diagnostic observation schedule-generic: a standard measure of social and communication deficits associated with the spectrum of autism. Journal of autism and developmental disorders. 30:205–223.

22. Constantino JN, Davis SA, Todd RD, Schindler MK, Gross MM, Brophy SL, et al. (2003): Validation of a brief quantitative measure of autistic traits: comparison of the social responsiveness scale with the autism diagnostic interview-revised. Journal of autism and developmental disorders. 33:427–433.

23. DuPaul GJ, Power TJ, Anastopoulos AD, Reid R (1998): ADHD Rating Scale—IV: Checklists, norms, and clinical interpretation. New York, NY, US: Guilford Press.

24. Moessnang C, Schafer A, Bilek E, Roux P, Otto K, Baumeister S, et al. (2016): Specificity, reliability and sensitivity of social brain responses during spontaneous mentalizing. Social cognitive and affective neuroscience. 11:1687–1697.

25. Plichta MM, Grimm O, Morgen K, Mier D, Sauer C, Haddad L, et al. (2014): Amygdala habituation: a reliable fMRI phenotype. NeuroImage. 103:383–390.

26. Damiano CR, Cockrell DC, Dunlap K, Hanna EK, Miller S, Bizzell J, et al. (2015): Neural mechanisms of negative reinforcement in children and adolescents with autism spectrum disorders. Journal of neurodevelopmental disorders. 7:12.

27. Schwarz K, Moessnang C, Schweiger JI, Baumeister S, Plichta MM, Brandeis D, et al. (2020): Transdiagnostic Prediction of Affective, Cognitive, and Social Function Through Brain Reward Anticipation in Schizophrenia, Bipolar Disorder, Major Depression, and Autism Spectrum Diagnoses. Schizophr Bull. 46:592–602.

28. Neuhaus E, Bernier RA, Beauchaine TP (2015): Electrodermal Response to Reward and Non-Reward Among Children With Autism. Autism research : official journal of the International Society for Autism Research. 8:357–370.

29. van Dongen EV, von Rhein D, O’Dwyer L, Franke B, Hartman CA, Heslenfeld DJ, et al. (2015): Distinct effects of ASD and ADHD symptoms on reward anticipation in participants with ADHD, their unaffected siblings and healthy controls: a cross-sectional study. Molecular autism. 6:48.

30. Greene RK, Spanos M, Alderman C, Walsh E, Bizzell J, Mosner MG, et al. (2018): The effects of intranasal oxytocin on reward circuitry responses in children with autism spectrum disorder. Journal of neurodevelopmental disorders. 10:12.

31. Wise RA (2004): Dopamine, learning and motivation. Nat Rev Neurosci. 5:483–494.

32. Green SA, Hernandez L, Bookheimer SY, Dapretto M (2016): Salience Network Connectivity in Autism Is Related to Brain and Behavioral Markers of Sensory Overresponsivity. J Am Acad Child Adolesc Psychiatry. 55:618–626 e611.

33. Uddin LQ, Supekar K, Lynch CJ, Khouzam A, Phillips J, Feinstein C, et al. (2013): Salience network-based classification and prediction of symptom severity in children with autism. JAMA psychiatry. 70:869–879.

34. Oldehinkel M, Mennes M, Marquand A, Charman T, Tillmann J, Ecker C, et al. (2019): Altered Connectivity Between Cerebellum, Visual, and Sensory-Motor Networks in Autism Spectrum Disorder: Results from the EU-AIMS Longitudinal European Autism Project. Biol Psychiatry Cogn Neurosci Neuroimaging. 4:260–270.

35. Neufeld J, Kuja-Halkola R, Mevel K, Cauvet E, Fransson P, Bolte S (2018): Alterations in resting state connectivity along the autism trait continuum: a twin study. Mol Psychiatry. 23:1659–1665.

36. Gerraty RT, Davidow JY, Foerde K, Galvan A, Bassett DS, Shohamy D (2018): Dynamic Flexibility in Striatal-Cortical Circuits Supports Reinforcement Learning. J Neurosci. 38:2442–2453.

37. van den Bos W, Cohen MX, Kahnt T, Crone EA (2012): Striatum-medial prefrontal cortex connectivity predicts developmental changes in reinforcement learning. Cerebral cortex. 22:1247–1255.

38. Gu R, Huang W, Camilleri J, Xu P, Wei P, Eickhoff SB, et al. (2019): Love is analogous to money in human brain: Coordinate-based and functional connectivity meta-analyses of social and monetary reward anticipation. Neurosci Biobehav Rev. 100:108–128.

39. Abrams DA, Padmanabhan A, Chen T, Odriozola P, Baker AE, Kochalka J, et al. (2019): Impaired voice processing in reward and salience circuits predicts social communication in children with autism. Elife. 8.

40. Abrams DA, Lynch CJ, Cheng KM, Phillips J, Supekar K, Ryali S, et al. (2013): Underconnectivity between voice-selective cortex and reward circuitry in children with autism. Proc Natl Acad Sci U S A. 110:12060–12065.

41. Bast N, Poustka L, Freitag CM (2018): The locus coeruleus-norepinephrine system as pacemaker of attention – a developmental mechanism of derailed attentional function in autism spectrum disorder. The European journal of neuroscience. 47:115–125.

42. von Rhein D, Cools R, Zwiers MP, van der Schaaf M, Franke B, Luman M, et al. (2015): Increased neural responses to reward in adolescents and young adults with attention-deficit/hyperactivity disorder and their unaffected siblings. J Am Acad Child Adolesc Psychiatry. 54:394–402.

43. Kappel V, Lorenz RC, Streifling M, Renneberg B, Lehmkuhl U, Strohle A, et al. (2015): Effect of brain structure and function on reward anticipation in children and adults with attention deficit hyperactivity disorder combined subtype. Social cognitive and affective neuroscience. 10:945–951.

44. Paloyelis Y, Mehta MA, Faraone SV, Asherson P, Kuntsi J (2012): Striatal sensitivity during reward processing in attention-deficit/hyperactivity disorder. J Am Acad Child Adolesc Psychiatry. 51:722–732 e729.

45. Bjork JM, Chen G, Smith AR, Hommer DW (2010): Incentive-elicited mesolimbic activation and externalizing symptomatology in adolescents. Journal of child psychology and psychiatry, and allied disciplines. 51:827–837.

46. Furukawa E, Bado P, Tripp G, Mattos P, Wickens JR, Bramati IE, et al. (2014): Abnormal striatal BOLD responses to reward anticipation and reward delivery in ADHD. PloS one. 9:e89129.

47. Wolfers T, Floris DL, Dinga R, van Rooij D, Isakoglou C, Kia SM, et al. (2019): From pattern classification to stratification: towards conceptualizing the heterogeneity of Autism Spectrum Disorder. Neurosci Biobehav Rev. 104:240–254.

