## Supplementary Material for "Attenuated Anticipation of Social and Monetary Rewards in Autism Spectrum Disorders"

### Sample description by age group

**Table S1:** Participant characteristics by sample and age group.

| **Children** | | | |
|---|---|---|---|
| | ASD | TD | group comparison |
| Total N | 38 | 40 | |
| Demographics | | | |
| Sex (m/f) | 29/9 | 24/16 | $\chi^2$(1)=2.382, *p*=.123 |
| Age (years) | 10.06 ± 1.35 (7.56 - 11.97) | 10.26 ± 1.26 (7.57 - 11.98) | *t*(76)=-.675, p=.502 |
| IQ (full IQ) | 112.55 ± 13.16 (86.36 - 148.00) | 110.42 ± 12.10 (76.00 - 133.00) | *t*(75)=.740, *p*=.462 |
| Handedness (right/left/ambidextrous/unknown) | 31/4/0/3 | 28/3/1/8 | $\chi^2$(3)=3.519, *p*=.318 |
| Medication use (no/yes/unknown) | 12/14/12 | 20/3/17 | $\chi^2$(2)=9.935, *p*=.007 |
| fMRI quality control | | | |
| SID Mean framewise displacement (FD; in mm) | .18 ± .08 (.05 - .36) | .14 ± .07 (.06 - .34) | *t*(76)=2.579, *p*=.012 |
| SID Volumes with FD >0.5 mm (in %) | 5.49 ± 6.08 (0 - 18.24) | 2.52 ± 4.10 (0 - 14.86) | *t*(64.51)=2.523, *p*=.014 |
| SID Signal-to-noise ratio | 11.18 ± .62 (9.96 - 12.51) | 11.20 ± .84 (8.28 - 13.10) | *t*(76)=-.130, *p*=.897 |
| MID Mean framewise displacement (FD; in mm) | .18 ± .07 (.05 - .29) | .16 ± .08 (.05 - .41) | *t*(76)=1.563, *p*=.122 |
| MID Volumes with FD >0.5 mm (in %) | 4.83 ± 4.59 (0 - 12.84) | 3.53 ± 4.58 (0 - 16.89) | *t*(76)=1.252, *p*=.215 |
| MID Signal-to-noise ratio | 11.35 ± .78 (10.05 - 13.43) | 11.60 ± 1.11 (8.22 - 14.08) | *t*(76)=-1.118, *p*=.267 |
| Clinical characteristics | | | |
| ADI-R | | | |
| Social interaction | 14.81 ± 6.77 (1 - 25) | | |
| Communication | 12.62 ± 5.93 (3- 24) | | |
| RRB | 3.97 ± 3.04 (0 - 12) | | |
| ADOS | | | |
| Social affect | 5.33 ± 2.32 (1 - 9) | | |
| RRB | 4.06 ± 2.87 (1 - 9) | | |
| Total | 4.58 ± 2.42 (1 - 9) | | |
| SRS-2 | | | |
| rawscore | 89.91 ±31.27 (32 - 163) | 19.68 ± 14.29 (2 - 74) | *t*(47.14)=11.813, *p*<.000 |
| t-score | 72.88 ± 11.98 (49- 90) | 45.12 ± 5.61 (37 - 66) | *t*(47.76)=12.129, *p*<.000 |
| ADHD reserach diagnosis*(ADHD/noADHD/missing) | 18/16/4 | 2/30/8 | $\chi^2$(1)=17.016, *p*<.001 |
| DAWBA comorbidities | | | |
| ADHD symptoms | 2.30 ± 1.51 (0 - 5) | .39 ± 1.03 (0 - 4) | *t*(50.45)=5.792 *p*<.000 |
| Anxiety symptoms | 2.77 ± 1.55 (0 - 5) | 1.21 ± .70 (0 - 4) | *t*(44.16)=5.464, *p*<.000 |
| Depression symptoms | .97 ± 1.33 (0-5) | .12 ± .33 (0-1) | *t*(34.94)=3.461, *p*=.001 |

| **Adolescents** | | | |
|---|---|---|---|
| | ASD | TD | *group comparison* |
| Total N | 88 | 61 | |
| Demographics | | | |
| Sex (m/f) | 68/20 | 39/22 | *χ2(1)=3.166, p=.075* |
| Age (years) | 15.09 ± 1.75 (12.07 - 17.90) | 15.61 ± 1.59 (12.44 - 17.99) | *t(147)=-1.841, p=.068* |
| IQ (full IQ) | 102.63 ± 15.19 (75.00 - 143.00) | 104.01 ± 12.46 (76.82 - 123.00) | *t(147)=-.584, p=.560* |
| Handedness (right/left/ambidextrous/unknown) | 60/10/4/14 | 45/5/0/11 | *χ2(3)=3.388, p=.336* |
| Medication use (no/yes/unknown) | 22/37/29 | 25/6/30 | *χ2(2)=18.264, p<.001* |
| fMRI quality control | | | |
| SID Mean framewise displacement (FD; in mm) | .13 ± .07 (.03 - .41) | .11 ± .06 (.05 - .29) | *t(147)=2.298, p=.023* |
| SID Volumes with FD >0.5 mm (in %) | 2.36 ± 3.47 (0 - 13.51) | 1.67 ± 3.16 (0 - 14.19) | *t(147)=1.228, p=.221* |
| SID Signal-to-noise ratio | 9.51 ± 1.23 (6.28 - 12.21) | 9.63 ± 1.11 (6.49 - 12.38) | *t(147)=-.596, p=.552* |
| MID Mean framewise displacement (FD; in mm) | .15 ± .08 (.03 - .36) | .13 ± .07 (.05 - .35) | *t(147)=2.106, p=.037* |
| MID Volumes with FD >0.5 mm (in %) | 3.62 ± 5.03 (0 - 19.59) | 2.23 ± 3.75 (0 - 16.22) | *t(147)=.671, p=.503* |
| MID Signal-to-noise ratio | 9.55 ± 1.33 (6.08 - 12.80) | 9.69 ± 1.23 (6.40 - 12.03) | *t(146.16)=1.942, p=.054* |
| Clinical characteristics | | | |
| ADI-R | | | |
| Social interaction | 16.85 ± 6.43 (2 - 29) | | |
| Communication | 13.21 ± 5.68 (1 - 26) | | |

| | | | |
|---|---|---|---|
| RRB | 3.99 ± 2.67 (0 - 12) | | |
| ADOS | | | |
| Social affect | 6.18 ± 2.72 (1 - 10) | | |
| RRB | 4.28 ± 2.32 (1 - 9) | | |
| Total | 5.36 ± 2.81 (1 - 10) | | |
| SRS-2 | | | |
| rawscore | 91.34 ±29.37 (22 - 151) | 22.40 ± 14.22 (1 - 67) | *t(115.98)=17.946, p<.000* |
| t-score | 73.03 ± 11.51 (45- 90) | 45.72 ± 5.35 (38 - 63) | *t(113.72)=18.318, p<.000* |
| ADHD reserach diagnosis*(ADHD/noADHD/missing) | 29/43/16 | 5/45/11 | *χ2(1)=13.457, p<.001* |
| DAWBA comorbidities | | | |
| ADHD symptoms | 2.03 ± 1.58 (0 - 5) | .18 ± .59 (0 - 3) | *t(87.43)=8.472, p<.000* |
| Anxiety symptoms | 2.52 ± 1.32 (0 - 5) | .73 ± .60 (0 - 2) | *t(119.87)=8.251, p<.000* |
| Depression symptoms | .80 ± 1.09 (0-4) | .35 ± .62 (0-2) | *t(118.00)=2.516, p=.013* |

| **Adults** | | | |
|---|---|---|---|
| | ASD | TD | *group comparison* |
| Total N | 86 | 80 | |
| Demographics | | | |
| Sex (m/f) | 60/26 | 52/28 | *χ2(1)=.429, p=.512* |
| Age (years) | 22.50 ± 3.49 (18.02 - 30.60) | 23.00 ± 3.16 (18.07 - 30.78) | *t(164)=-.966, p=.336* |
| IQ (full IQ) | 105.94 ± 14.36 (75.56 - 148.00) | 108.40 ± 12.30 (75.56 - 141.00) | *t(164)=-1.179, p=.240* |
| Handedness (right/left/ambidextrous/unknown) | 58/12/4/12 | 49/7/3/21 | *χ2(3)=4.459, p=.216* |
| Medication use (no/yes/unknown) | 30/31/25 | 27/3/50 | *χ2(2)=31.374, p<.001* |
| fMRI quality control | | | |
| SID Mean framewise displacement (FD; in mm) | .09 ± .05 (.03 - .27) | .09 ± .06 (.03 - .34) | *t(164)=-.115, p=.908* |
| SID Volumes with FD >0.5 mm (in %) | .95 ± 2.00 (0 - 11.49) | 1.23 ± 3.00 (0 - 16.22) | *t(164)=-.717, p=.474* |
| SID Signal-to-noise ratio | 9.40 ± 1.03 (7.10 - 11.97) | 9.45 ± 1.02 (6.93 - 12.13) | *t(164)=-.328, p=.744* |
| MID Mean framewise displacement (FD; in mm) | .10 ± .06 (.03 - .31) | .10 ± .06 (.03 - .36) | *t(164)=-.195, p=.845* |
| MID Volumes with FD >0.5 mm (in %) | 1.13 ± 2.65 (0 - 15.54) | 1.21 ± 2.46 (0 - 15.54) | *t(164)=-.192, p=.848* |
| MID Signal-to-noise ratio | 9.45 ± 1.17 (7.05 - 13.62) | 9.44 ± 1.06 (6.82 - 11.77) | *t(164)=.015, p=.988* |
| Clinical characteristics | | | |
| ADI-R | | | |
| Social interaction | 14.27 ± 6.48 (0 - 28) | | |
| Communication | 11.63 ± 5.54 (0 - 24) | | |
| RRB | 3.84 ± 2.52 (0 - 12) | | |
| ADOS | | | |
| Social affect | 5.46 ± 2.40 (1 - 10) | | |
| RRB | 4.39 ± 2.38 (1 - 10) | | |
| Total | 4.51 ± 2.42 (1 - 10) | | |
| SRS-2 | | | |
| rawscore | 76.06 ±29.34 (20 - 143) | 29.27 ± 15.07 (4 - 87) | *t(123.83)=12.270, p<.000* |
| t-score | 62.56 ± 10.33 (43- 86) | 46.15 ± 5.33 (37 - 66) | *t(124.09)=12.215, p<.000* |
| ADHD reserach diagnosis*(ADHD/noADHD/missing) | 69/118/25 | 11/130/40 | *χ2(1)=9.376, p=.002* |
| DAWBA comorbidities | | | |
| ADHD symptoms | 1.20 ± 1.41 (0 - 4) | | |
| Anxiety symptoms | 2.59 ± 1.14 (0 - 4) | | |
| Depression symptoms | .82 ± 1.29 (0-5) | | |

Participant characteristics, split by age group. ADI-R: Autism Diagnostic Interview-Revised. Scores were computed for reciprocal interaction (social interaction), communication, and restrictive, repetitive stereotyped behaviors and interests (RRB). ADOS-2: Autism Diagnostic Observation Schedule 2. Calibrated severity scores were computed for social affect, restricted and repetitive behaviors (RRB) and the overall total score. SRS-2: Social Responsiveness Scale-2. Total raw and total T scores (sex+age normalized) are reported. The raw SRS-2 scores were used in our analyses. ADHD research diagnosis was based on applying DSM-V criteria to symptom scores in the parent- and self-rated ADHD rating scale. Self-rated scores were used when parent-rated scores were not available. Comorbid symptoms of ADHD, depression and anxiety were assessed with the Development and Well Being Assessment (DAWBA), generating six levels (ordinal scores 0 to 5) of prediction of the probability of a disorder (~0.1%, ~0.5%, ~3%, ~15%, ~50%, >70%). SID social incentive delay task, MID monetary incentive delay task.

### Standard operation procedures and quality control

Standard operation procedures were implemented to harmonize data acquisition between sites and across time (1). This included hands-on and face-to-face training according to detailed protocols before study rollout and regular exchange between sites during data acquisition. Hard- and software for data acquisition was aligned as closely as possible between sites. Procedures were undertaken to optimize the MRI sequences for the best scanner-specific options while harmonizing across sites, and phantoms and travelling heads were employed to assure standardization and quality assurance of the multi-site image-acquisition. Test-retest reliability of the fMRI task battery was ensured (2-4).

Of the total sample, n=285 ASD participants and n=217 typically developing (TD) individuals had an IQ above 75 and data for both the MID and SID available. Several quality assessment (QA) metrics were calculated (http://preprocessed-connectomes-project.org/quality-assessment-protocol/) for these participant's datasets. Head motion was quantified as frame-wise displacement (FD, (5)), with two separate scores extracted for each dataset: mean FD and percent of volumes exceeding 0.5 mm FD. Additional temporal QA metrics comprised the temporal signal-to-noise ratio and the average change of a volume's mean intensity across time points (DVARS (6)). We additionally calculated the signal-to-noise ratio (SNR) based on the mean volume of the realigned time-series. In the present study, bad functional data quality was defined as excessive head movement with more than 20 percent of frames with a frame-wise displacement (FD) > 0.5mm (5) and/or signal loss with less than 80% overlap of the individual brain mask with the template mask in MNI space.

Participants were excluded based on anatomical brain abnormality (n=16), incomplete fMRI scan (n=12), technical problems (n=77), incorrect task performance (n=21), bad functional data quality (N=111) and data corruption (n=4) or a combination of these reasons. A comparison of the main QA metrics between ASD and TD in the final sample is presented in table 1 in the main text.

### Experimental paradigm

Participants were asked to give a speeded response (button press) to a visual target screenflash. A cue arrow pointing upwards indicated the possibility to obtain a reward if responses were given within a predefined response time window (win trial). No reward option was given in trials preceded by a horizontal cue arrow (neutral trial). The response time window was continuously adapted to ensure a comparable number of reward events across subjects and groups (~60 %). Sufficiently fast responses on win trials were followed by the presentation of a 2€/2£ coin in the MID and a smiling female face in the SID as feedback. Blurred control stimuli were presented in neutral trials and as feedback following slow responses in win trials. In total, 30 win trials and 30 neutral trials were presented in a pseudorandomized order during each task. SID and MID were collected as separate paradigms with SID always presented first, followed by MID. Task order was not randomized across participants in order to avoid loss of motivation after performing the monetary task. Note that the feedback presentation was temporally decoupled from the target presentation but not from the button press. For a visualization of the task design and stimuli, please refer to figure 1 in the main text.

### First level fMRI data analysis

SID and MID tasks were combined as two sessions in a general linear model (GLM) on the single subject level. Each session was modeled using the six task regressors (cue win, cue neutral, target, feedback win, feedback lost and feedback neutral) and six realignment parameters were included for both tasks. All regressors were modeled as stick functions, convolved with a canonical hemodynamic response function (HRF). At the model estimation stage, a high-pass filter with a cutoff of 128 seconds and an autoregressive model of the first order were applied. Contrast images (cue win > cue neutral; feedback won > feedback neutral) were created for each task separately and for both tasks combined. Additionally, a contrast image for the interaction between condition (win, neutral) and task (SID, MID) was calculated.

### Behavioural data

Behaviourally, individuals with autism have been reported to show decreased accuracy compared to typically developing (TD) individuals in these tasks in some cases (7, 8) but not others (9-11). Typically, reaction times (RT) are faster when a subsequent reward is possible for all participants and do not differ between groups (7, 9, 10, 12, 13), but see (11, 14).

Reaction times (RT) and accuracy (percentage of successful trials) were analyzed using SPSS Software package (Version 25, IBM Corp., Armonk, NY, USA). A repeated measures ANOVAs with the within subject factors condition (win, neutral) and task (MID, SID) and between subject factor of diagnosis (TD, ASD) and covariates age (mean-centered) sex, and study sites (dummy coded) were used to assess effects of task, condition and diagnosis.

There were no significant effects of diagnosis, task or condition on RT. Accuracy was higher for win ($M$=70.5) compared to neutral trials ($M$=49.9, $F_{(1,380)}$=25.989, $p$<.001, $\eta^2$=.064). A significant task*condition interaction ($F_{(1,380)}$=3.896, $p$=.049, $\eta^2$=.010) revealed higher accuracy during win trials in the MID ($M$=73.2) compared to the SID ($M$=67.8, $p$<.001) while during neutral trials accuracy was higher in the SID ($M$=51.6) compared to the MID ($M$=48.2, $p$<.001). There was no significant effect of diagnosis on accuracy.

### Whole brain activation analyses for tasks separately

Although there was a significant interaction between task and condition during both reward anticipation and delivery, revealing stronger differential activation in the MID compared to the SID (see fig 2 and table 2 in main text), this interaction was not different between individuals with ASD and TD and not associated with SRS-2 scores. However, here we still assess differences between ASD and TD in the SID and MID separately (see figures S1 and S2, and tables S2 and S3).

The separate analysis of the MID revealed similar results to the combined analysis, with the effect of diagnostic group remaining significant at the whole brain level in the right VS ($F_{(1,384)}$=22.53, $p_{FWE}$=.022, k=9). In contrast, separate analysis of the SID revealed no significant effect of diagnosis at the whole brain level. For details, see figure S1 and tables S2 and S3. Explorative ROI analyses in the SID and MID separately revealed that group differences remained significant in the MID (left VS: $F_{(1,384)}$=15.201, $p$<.001, partial $\eta^2$=.038, right VS $F_{(1,384)}$=19.462, $p$<.001, partial $\eta^2$=.048) and in the right VS for the SID ($F_{(1,384)}$=6.732, $p$=.010, partial $\eta^2$=.017). Effects in the left VS for the SID did not survive correction for multiple testing ($F_{(1,384)}$=4.876, $p$=.028, partial $\eta^2$=.013). For details, see figure S1 and supplementary table S3.

Separate analyses of the MID and SID also yielded no significant effect of diagnosis in either task during reward delivery. For details, see figure S2 and supplementary tables S2 and S3). Explorative ROI analysis in the MID revealed that the effect of diagnosis reached significance in the right VS: ($F_{(1,370)}$=5.557, $p$=.019, partial $\eta^2$=.015) while it remained below threshold for the left VS ($F_{(1,370)}$=4.804, $p$=.029, partial $\eta^2$=.013). There was no significant effect of diagnosis in the SID (left VS: $F_{(1,370)}$=1.383, $p$=.240, partial $\eta^2$=.004, right VS $F_{(1,370)}$=.802, $p$=.371, partial $\eta^2$=.002). For details, see figure S2 and table S3.

Across both tasks, there was also no effect of continuous SRS-2 scores on the whole brain or ROI level.

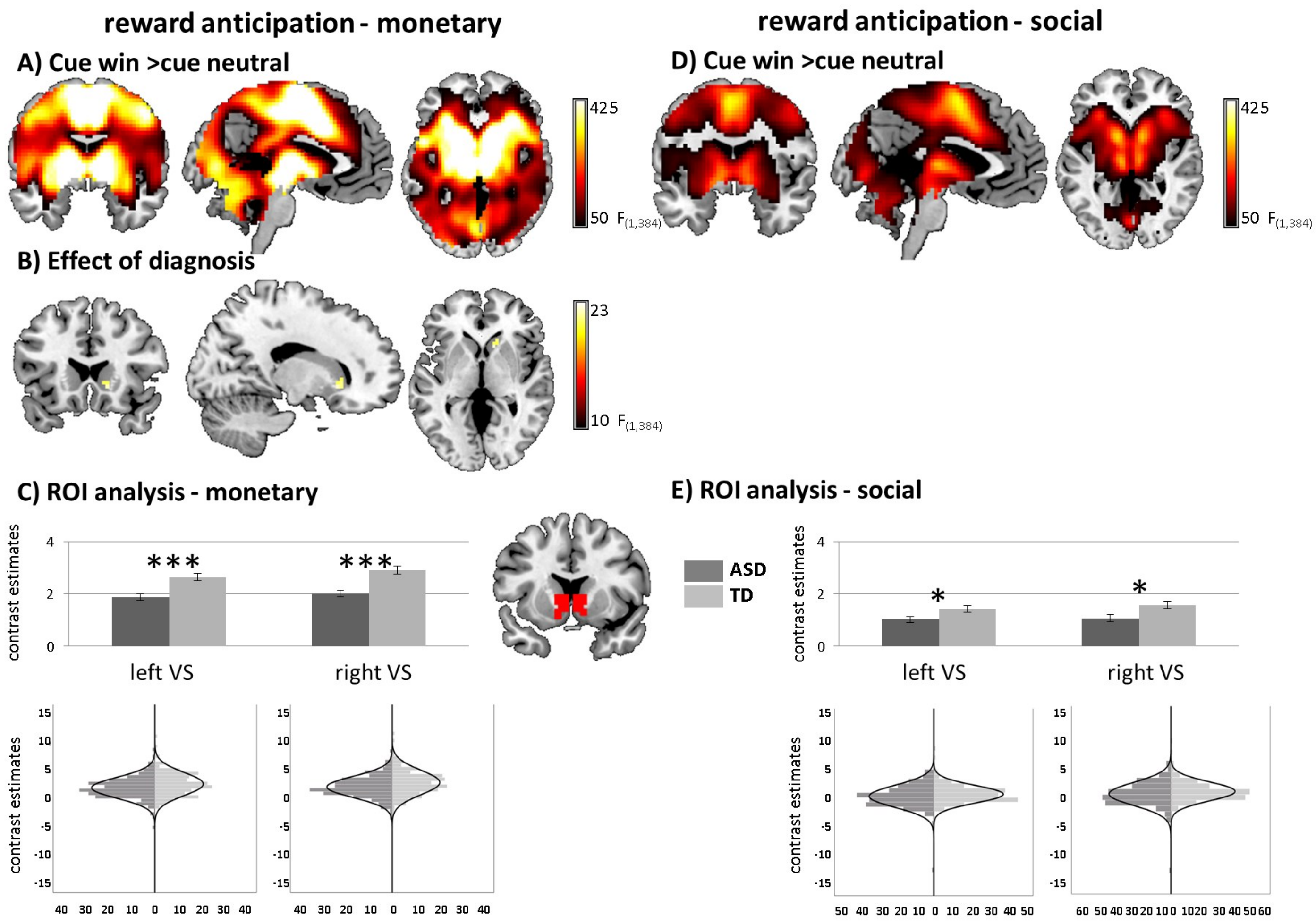

**Figure S1**: Whole-brain familywise error corrected brain activation to win compared to neutral cues for the MID and SID separately. A) activation across both ASD and TD individuals in the MID. B) Effect of diagnosis in right ventral striatum in the MID. C) Effect of diagnosis in the region of interest (ROI) analysis of the left and right ventral striatum with corresponding distribution plots for the MID. D) activation across both ASD and TD individuals in the SID. E) Effect of diagnosis in the region of interest (ROI) analysis of the left and right ventral striatum with corresponding distribution plots for the SID. ****p*<.001, **p*<.05.

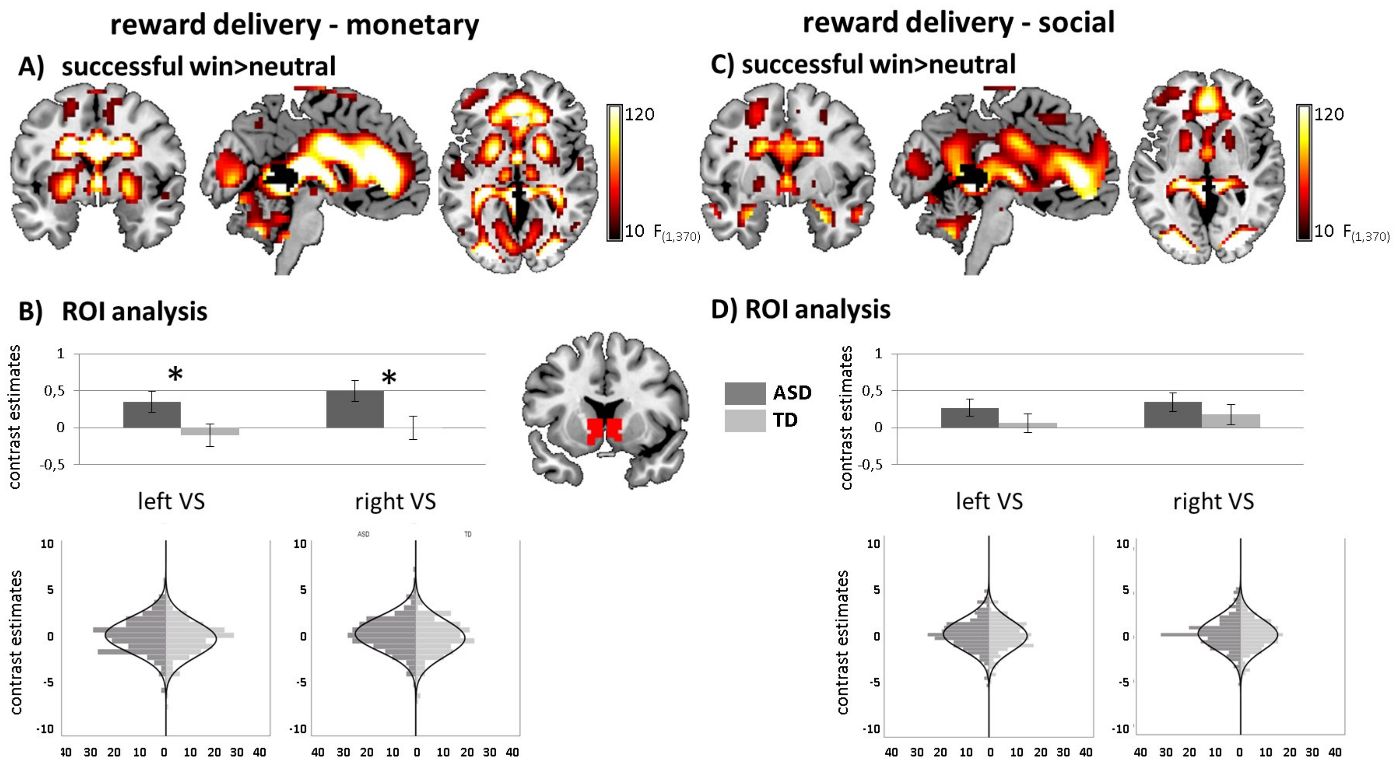

**Figure S2**: Whole-brain familywise error corrected brain activation to successful win compared to neutral trials for the MID and SID separately. A) activation across both ASD and TD individuals in the MID. B) Effect of diagnosis in the region of interest (ROI) analysis of the left and right ventral striatum with corresponding distribution plots for the MID. C) activation across both ASD and TD individuals in the SID. D) Effect of diagnosis in the region of interest (ROI) analysis of the left and right ventral striatum with corresponding distribution plots for the SID. **p*<.05.

**Table S2:** Whole-brain effects of brain activation for SID and MID separately.

| Region | Hemisphere | Direction | k | x | y | z | F | $p_{(FWE\text{-}corr)}$ |
|---|---|---|---|---|---|---|---|---|
| **ANTICIPATION MID** | | | | | | | | |
| **EFFECT OF TASK** | | | | | | | | |
| supplementary motor area | r | win>neutral | 42326 | 3 | 2 | 53 | 870.267 | 0.000 |
| nucleus accumbens | r | win>neutral | | 12 | 8 | -4 | 815.828 | 0.000 |
| supplementary motor area | l | win>neutral | | -6 | 8 | 44 | 797.693 | 0.000 |
| pallidum | l | win>neutral | | -12 | 8 | -4 | 793.943 | 0.000 |
| middle cingulate gyrus | r | win>neutral | | 9 | 11 | 41 | 781.462 | 0.000 |
| thalamus, intralaminar | l | win>neutral | | -9 | -19 | -1 | 776.746 | 0.000 |
| thalamus, mediodorsal lateral parvocellular | r | win>neutral | | 9 | -16 | -1 | 707.410 | 0.000 |
| precentral gyrus | l | win>neutral | | -36 | -16 | 50 | 655.909 | 0.000 |
| precentral gyrus | l | win>neutral | | -36 | -10 | 53 | 655.081 | 0.000 |
| precentral gyrus | l | win>neutral | | -27 | -28 | 65 | 626.463 | 0.000 |
| middle cingulate gyrus | l | win>neutral | | -9 | -25 | 44 | 563.826 | 0.000 |
| insula | r | win>neutral | | 30 | 26 | 2 | 533.024 | 0.000 |
| insula | l | win>neutral | | -27 | 23 | 5 | 530.611 | 0.000 |
| precentral gyrus | l | win>neutral | | -24 | -10 | 65 | 517.364 | 0.000 |
| middle cingulate gyrus | r | win>neutral | | 9 | -28 | 47 | 512.844 | 0.000 |

| | | | | | | | | |
|---|---|---|---|---|---|---|---|---|
| middle frontal gyrus | r | win>neutral | | 39 | -7 | 53 | 482.072 | 0.000 |
| **EFFECT OF DIAGNOSIS** | | | | | | | | |
| caudate | r | TD>ASD | 9 | 12 | 17 | -1 | 22.533 | 0.022 |
| **DELIVERY MID** | | | | | | | | |
| **EFFECT OF TASK** | | | | | | | | |
| middle occipital gyrus | r | s. win>neutral | 9359 | 30 | -88 | -1 | 744.741 | 0.000 |
| middle occipital gyrus | l | s. win>neutral | | -24 | -94 | -1 | 734.134 | 0.000 |
| inferior occipita gyrus | l | s. win>neutral | | -27 | -88 | -7 | 637.429 | 0.000 |
| pallidum | r | neutral>s. win | 420 | 21 | 5 | -1 | 137.279 | 0.000 |
| insula | r | neutral>s. win | | 36 | -7 | 8 | 33.433 | 0.000 |
| insula | l | s. win>neutral | 165 | -33 | 14 | -16 | 132.034 | 0.000 |
| pallidum | l | neutral>s. win | 469 | -21 | 2 | -1 | 123.290 | 0.000 |
| insula | l | | | -36 | -7 | 5 | 36.622 | 0.000 |
| superior temporal gyrus | l | | | -39 | -13 | -7 | 21.866 | 0.032 |
| angular gyrus | r | neutral>s. win | 694 | 60 | -55 | 32 | 98.575 | 0.000 |
| middle temporal gyrus | r | neutral>s. win | | 54 | -64 | 11 | 93.408 | 0.000 |
| supramarginal gyrus | r | neutral>s. win | | 66 | -19 | 32 | 79.452 | 0.000 |
| lingual gyrus | l | neutral>s. win | 2393 | -6 | -79 | 2 | 90.596 | 0.000 |
| calcarine | r | neutral>s. win | | 6 | -79 | 5 | 87.877 | 0.000 |
| angular gyrus | l | | | -48 | -64 | 47 | 75.276 | 0.000 |
| middle temporal gyrus | l | neutral>s. win | 183 | -45 | -70 | 8 | 77.507 | 0.000 |
| vermis | | neutral>s. win | 34 | 0 | -37 | -40 | 75.909 | 0.000 |
| middle frontal gyrus | l | neutral>s. win | 967 | -39 | 20 | 47 | 74.903 | 0.000 |
| middle frontal gyrus | l | neutral>s. win | | -36 | 26 | 41 | 66.322 | 0.000 |
| middle frontal gyrus | l | neutral>s. win | | -36 | 38 | 26 | 57.545 | 0.000 |
| supplementary motor area | l | s. win>neutral | 70 | 0 | -13 | 74 | 74.385 | 0.000 |
| supplementary motor area | l | s. win>neutral | | 0 | 2 | 71 | 50.649 | 0.000 |
| paracentral lobule | r | s. win>neutral | | 0 | -28 | 74 | 49.733 | 0.000 |
| superior frontal gyrus | l | neutral>s. win | 145 | -21 | -7 | 59 | 59.315 | 0.000 |
| superior frontal gyrus | r | neutral>s. win | 300 | 27 | 50 | 20 | 44.435 | 0.000 |
| middle frontal gyrus | r | neutral>s. win | | 30 | 44 | 26 | 40.945 | 0.000 |
| middle frontal gyrus | r | neutral>s. win | | 36 | 23 | 38 | 37.056 | 0.000 |
| supplementary motor area | r | neutral>s. win | 126 | 12 | 5 | 47 | 38.244 | 0.000 |
| superior frontal gyrus | r | neutral>s. win | | 18 | -4 | 65 | 33.302 | 0.000 |
| paracentral lobule | r | | | 9 | -28 | 62 | 26.585 | 0.004 |
| rolandic operculum | r | neutral>s. win | 49 | 60 | 8 | 14 | 36.222 | 0.000 |
| precentral gyrus | l | s. win>neutral | 12 | -24 | -22 | 74 | 35.440 | 0.000 |
| precentral gyrus | r | neutral>s. win | 33 | 30 | -7 | 50 | 33.639 | 0.000 |
| Inferior frontal gyrus, opercular part | l | neutral>s. win | 31 | -54 | 5 | 11 | 31.498 | 0.001 |
| superior temporal gyrus | r | neutral>s. win | 50 | 51 | 2 | -13 | 28.538 | 0.002 |
| superior temporal gyrus | r | neutral>s. win | | 54 | -7 | -1 | 28.514 | 0.002 |

| | | | | | | | | |
|---|---|---|---|---|---|---|---|---|
| Heschl's gyrus | r | neutral>s. win | 8 | 39 | -28 | 14 | 25.125 | 0.008 |
| Inferior frontal gyrus, triangular part | r | neutral>s. win | 5 | 57 | 26 | 14 | 22.722 | 0.022 |
| **ANTICIPATION SID** | | | | | | | | |
| **EFFECT OF TASK** | | | | | | | | |
| putamen | l | win>neutral | 35668 | -15 | 5 | -7 | 366.221 | 0.000 |
| nucleus accumbens | r | win>neutral | | 12 | 8 | -7 | 347.367 | 0.000 |
| supplementary motor area | l | win>neutral | | -3 | -1 | 59 | 338.159 | 0.000 |
| precentral gyrus | l | win>neutral | | -42 | -13 | 53 | 329.747 | 0.000 |
| thalamus, mediodorsal medial magnocellular | l | win>neutral | | -6 | -19 | 8 | 320.730 | 0.000 |
| supplementary motor area | r | win>neutral | | 6 | 5 | 50 | 301.722 | 0.000 |
| thalamus, mediodorsal medial magnocellular | r | win>neutral | | 6 | -13 | 5 | 297.261 | 0.000 |
| middle cingulate gyrus | l | win>neutral | | -6 | 11 | 38 | 287.338 | 0.000 |
| red nucleus | l | win>neutral | | -9 | -19 | -7 | 274.031 | 0.000 |
| red nucleus | l | win>neutral | | -6 | -22 | -10 | 271.665 | 0.000 |
| precentral gyrus | l | win>neutral | | -27 | -28 | 59 | 262.482 | 0.000 |
| red nucleus | r | win>neutral | | 9 | -19 | -10 | 259.582 | 0.000 |
| middle frontal gyrus | r | win>neutral | | 42 | -7 | 56 | 231.990 | 0.000 |
| insula | l | win>neutral | | -30 | 26 | 2 | 214.125 | 0.000 |
| lingual gyrus | l | win>neutral | | 0 | -76 | 2 | 207.940 | 0.000 |
| insula | r | win>neutral | | 33 | 26 | -4 | 202.890 | 0.000 |
| **DELIVERY SID** | | | | | | | | |
| **EFFECT OF TASK** | | | | | | | | |
| Calcarine fissure and surrounding cortex | r | s. win>neutral | 8891 | 24 | -91 | -1 | 907.615 | 0.000 |
| inferior occipita gyrus | l | s. win>neutral | | -24 | -91 | -4 | 695.419 | 0.000 |
| middle occipital gyrus | l | s. win>neutral | | -21 | -97 | 5 | 645.510 | 0.000 |
| middle frontal gyrus | l | neutral>s. win | 1877 | -39 | 35 | 29 | 89.398 | 0.000 |
| middle frontal gyrus | l | neutral>s. win | | -33 | 44 | 14 | 63.507 | 0.000 |
| superior frontal gyrus | l | neutral>s. win | | -21 | 2 | 53 | 59.495 | 0.000 |
| inferior parietal gyrus | l | neutral>s. win | 1153 | -36 | -43 | 44 | 79.507 | 0.000 |
| inferior parietal gyrus | l | neutral>s. win | | -51 | -28 | 38 | 72.565 | 0.000 |
| precuneus | l | neutral>s. win | | -12 | -67 | 53 | 69.840 | 0.000 |
| supramarginal gyrus | r | neutral>s. win | 303 | 63 | -22 | 35 | 63.666 | 0.000 |
| supramarginal gyrus | r | neutral>s. win | | 36 | -37 | 44 | 44.921 | 0.000 |
| supramarginal gyrus | r | | | 57 | -31 | 47 | 32.139 | 0.000 |
| cuneus | r | neutral>s. win | 438 | 6 | -85 | 17 | 57.019 | 0.000 |
| Inferior frontal gyrus, opercular part | r | neutral>s. win | 174 | 57 | 11 | 11 | 56.174 | 0.000 |
| insula | r | neutral>s. win | | 39 | 20 | 5 | 30.799 | 0.001 |
| superior parietal gyrus | r | | 109 | 18 | -67 | 53 | 54.793 | 0.000 |
| middle frontal gyrus | r | neutral>s. win | 311 | 36 | 32 | 35 | 54.568 | 0.000 |
| superior frontal gyrus | r | neutral>s. win | | 27 | 50 | 14 | 33.088 | 0.000 |
| supplementary motor area | r | s. win>neutral | 34 | 3 | -13 | 74 | 51.326 | 0.000 |

| | | | | | | | | |
|---|---|---|---|---|---|---|---|---|
| pallidum | r | neutral>s. win | 71 | 18 | 8 | -1 | 35.084 | 0.000 |
| superior frontal gyrus | r | | 28 | 27 | -7 | 53 | 27.116 | 0.004 |
| angular gyrus | l | s. win>neutral | 20 | -45 | -64 | 23 | 26.356 | 0.005 |
| middle temporal gyrus | | s. win>neutral | 8 | -51 | -40 | 2 | 23.989 | 0.015 |

Table provides test statistic of significant peak voxel(s) for whole-brain analysis. Voxel-level statistics were family-wise error (FWE) corrected for the number of voxels across the whole brain for each test. Significance was defined as pFWE<.05 with a cluster threshold of k≥5. Significant whole-brain results are localized in MNI space and labeled according to the automated anatomical labeling atlas 3 (aal3). SID social incentive delay task, MID monetary incentive delay task, ASD autism spectrum disorder, TD typically developing, s. win successful win.

**Table S3**: Overview over effects of diagnosis (categorical) and autism traits (dimensional) on functional brain activation separately in the MID and SID task.

| | TD vs. ASD | | | effect of SRS-2 | | |
|---|---|---|---|---|---|---|
| | whole-brain | left VS | right VS | whole-brain | left VS | right VS |
| original results (reported in main text) | | | | | | |
| anticipation | *right VS* $F_{(1,384)}$=22.84, $p_{FWE}$=.017 | $F_{(1,384)}$=14.163, *p*<.001 | $F_{(1,384)}$=18.693, *p*<.001 | no effect | no effect | no effect |
| delivery | no effect | $F_{(1,370)}$=4.829, *p*=.029 | $F_{(1,370)}$=4.719, *p*=.030 | no effect | no effect | no effect |
| MID | | | | | | |
| anticipation | right VS: $F_{(1,384)}$=22.53, $p_{FWE}$=.022 | $F_{(1,384)}$=15.20, *p*<.001 | $F_{(1,384)}$=19.46, *p*<.001 | no effect | no effect | no effect |
| delivery | no effect | $F_{(1,370)}$=4.80, *p*=.029 | **$F_{(1,370)}$=5.56, *p*=.019** | no effect | no effect | no effect |
| SID | | | | | | |
| anticipation | **no effect** | $F_{(1, 384)}$=4.88, *p*=.028 | $F_{(1, 384)}$=6.73, *p*=.010 | no effect | no effect | no effect |
| delivery | no effect | no effect | no effect | no effect | no effect | no effect |

Table provides test statistic of region of interest (ROI) analysis and of significant peak voxel(s) for whole-brain analysis. Voxel-level statistics were family-wise error (FWE) corrected for the number of voxels across the whole brain for each test. Significance was defined as pFWE<.05 with a cluster threshold of k≥5. To correct for investigating left and right VS activity separately in the ROI analysis, critical alpha was adjusted to p<.025 based on the Bonferroni procedure. Deviations from results reported in the main text are highlighted in bold. Significant whole-brain results are localized in MNI space and labeled according to the automated anatomical labeling atlas (aal). Abbreviations: TD typically developing, ASD autism spectrum disorder, VS ventral striatum, SID social incentive delay task, MID monetary incentive delay task, IA interaction.

### Control analyses

Please note that we did not correct for the number of tests performed in the control analyses to maximize sensitivity.

### Age group

Previous research in TD individuals has demonstrated several, but inconsistent age effects (for reviews, see 15, 16). Regarding reward anticipation, studies have reported hypoactivation in adolescents compared to adults and children (17-19). Regarding reward consumption or responses to entire reward processing trials, studies reported either no age-related effects (18, 20) or increased activity in adolescence in the VS (21-24) or dorsal caudate (25). Other studies show a linear increase of VS activation with age during reward anticipation (26), a linear decrease during reward delivery (26) or a linear increase of VS activity assessing the entire trial (27). Further, the pattern of age-related differences in reward processing in ASD remains elusive. We hypothesize a similar quadratic effect of age on VS activity in both TD and ASD individuals. We assessed linear and quadratic effects of age as well as interactions between age and diagnosis dimensionally across the whole sample in separate models where age and $age^2$ were added as additional covariates of interest.

During reward anticipation, no linear or non-linear effects of age were observed.

During reward delivery, a linear increase of activation with increasing age was observed in the right superior medial frontal gyrus ($F_{(1,370)}$=23.58, $p_{FWE}$=.016, $k$=8, figure S3 A). There was no linear association of age and VS. Quadratic age yielded an effect on the whole-brain level in a cluster comprising the left amygdala and pallidum ($F_{(1,369)}$=27.10, $p_{FWE}$=.004, $k$=18, figure S3 B). A trend for a quadratic effect of age was observed in the right VS ($F_{(1,367)}$=3.609, $p$=.058) and is illustrated in figure S3 C. In the left VS a trend for an interaction between $age^2$ and reward type was observed ($F_{(1,367)}$=3.626, $p$=.058) and is illustrated in figure S3 C. Follow up analysis revealed a significant effect of $age^2$ in the MID ($F_{(1,368)}$=5.242, $p$=.023), but not in the SID ($F_{(1,368)}$=.010, $p$=.921). No interactions between age or $age^2$ and diagnosis were observed.

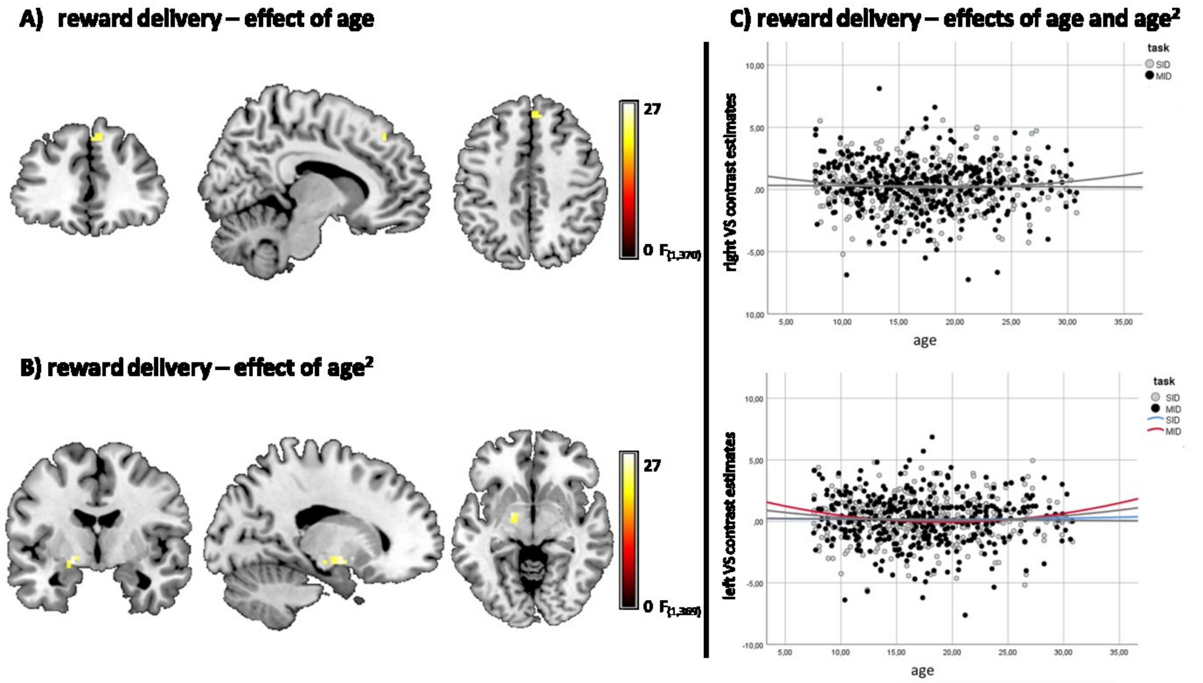

**Figure S3**: Linear and non-linear effects of age. A) Whole-brain familywise error corrected brain activation associated with age for the contrast of won compared to neutral trials. B) Whole-brain familywise error corrected brain activation associated with age$^2$ for the contrast of won compared to neutral trials. C) Scatterplot with linear and quadratic model fits for the association between age and contrast estimates of right and left ventral striatal activation across social and monetary reward for the contrast of won compared to neutral trials. Plot for left VS includes quadratic model fits for both tasks separately (MID in red, SID in blue).

Investigating effects of age yielded a linear increase of activity within the right superior medial frontal gyrus during reward delivery. This region in the medial prefrontal cortex (mPFC) has been shown to encode task value and expectation of monetary reward (28) and more frontal parts of this region have been associated with valuation of social interactions (29, 30). Further, our results are in line with previous studies reporting increased activation in the mPFC in older compared to younger participants during reward delivery (27, 31, 32). The trend-level significant effects for age$^2$ and the interaction between task and age$^2$ in the right VS and left VS, respectively, revealed lowest VS activity in adolescence and early adulthood with higher activation in childhood and towards the upper end of the age range. Importantly, our results are in contrast to findings of Schreuders et al. (24) showing a quadratic association of VS activity during delivery and age, with peak VS activity levels in adolescence. While this study was similar in sample size and age-range, the diverging results may be explained by the different tasks applied and, more importantly, by the fact that Schreuders et al. employed a longitudinal design across three assessment timepoints. The longitudinal approach of the LEAP study will allow us to test the replicability of our finding in future investigations. Further, as none of the age-related effects we observed differed significantly between individuals with ASD and TD, a within-subject design should complement the current cross-sectional analyses with higher sensitivity for differences in developmental changes of motivational processes due to a better control of between- and within-subject confounds.

To allow comparability to previous studies explore effects within narrower age-ranges, we repeated these main analyses within age-specific subgroups: children (7.0-11.9 years), adolescents (12.0-17.9 years) and adults (18.0-30.9 years).

A main effect of diagnosis was found only in the adult subsample (see table S4 and figure S4 A). However, in children a significant interaction between task and diagnosis emerged in the right VS during reward anticipation with increased VS activation in TD compared to ASD only during the MID ($p$=.003) but not during the SID task ($p$=.896). During reward delivery, the interaction between task and diagnosis in the right VS was significant at trend-level in the childhood subsample with increased right VS activation to monetary compared to social rewards in ASD ($p$=.022), but not in TD children ($p$=.562 ). For details see table S4 and figure S4 B.

**Table S4**: Overview over effects of diagnosis and age on functional brain activation for adults, adolescents and children separately.

| | TD vs. ASD | | | effect of age | | | quadratic effect of age | | | effect of SRS-2 | | |
|---|---|---|---|---|---|---|---|---|---|---|---|---|
| | whole-brain | left VS | right VS | whole-brain | left VS | right VS | whole-brain | left VS | right VS | whole-brain | left VS | right VS |
| full sample (group comparison as reported in main text) | | | | | | | | | | | | |
| anticipation | *right VS*<br>$F_{(1,384)}$=22.84, $p_{FWE}$=.017 | $F_{(1,384)}$=14.163, *p*<.001 | $F_{(1,384)}$=18.693, *p*<.001 | no effect | no effect | no effect | no effect | no effect | no effect | no effect | no effect | no effect |
| delivery | no effect | $F_{(1,370)}$=4.829, *p*=.029 | $F_{(1,370)}$=4.719, *p*=.030 | *right superior medial frontal gyrus*<br>$F_{(1,370)}$=23.58, $p_{FWE}$=.016 | no effect | no effect | *left amygdala/pallidum*:<br>$F_{(1,367)}$=27.10, $p_{FWE}$=.004;<br>*left hippo-campus*:<br>$F_{(1,367)}$=22.18, $p_{FWE}$=.029 | no effect<br>*IA task*age_quad*<br>$F_{(1,367)}$=3.626, *p*=.058 | ($F_{(1,367)}$=3.609, *p*=.058 | no effect | no effect | no effect |
| adults (n=166) | | | | | | | | | | | | |
| anticipation | ***Superior parietal lobule:***<br>$\mathbf{F_{(1,384)}}$**=23.08, $\mathbf{p_{FWE}}$=.028** | $F_{(1,157)}$=10.561, *p*=.001 | $F_{(1,157)}$=11.504, *p*=.001 | no effect | no effect | no effect | no effect | no effect | no effect | no effect | no effect | no effect |
| delivery | no effect | $F_{(1,152)}$=11.186, *p*=.001 | $\mathbf{F_{(1,152)}}$**=10.631, *p*=.001** | **no effect** | no effect | no effect | **no effect** | no effect | no effect | no effect | no effect | no effect |
| adolescents (n=149) | | | | | | | | | | | | |
| anticipation | **no effect** | **no effect** | **no effect** | no effect | no effect | no effect | no effect | no effect | no effect | no effect | no effect | no effect |
| delivery | no effect | **no effect** | **no effect** | **no effect** | no effect | no effect | **no effect** | no effect | no effect | no effect | no effect | no effect |
| children (n=78) | | | | | | | | | | | | |
| anticipation | **no effect** | **no effect** | **no effect**<br>*IA task*diagnosis*<br>$F_{(1,70)}$=5.451, *p*=.022 | no effect | no effect | no effect | no effect | no effect | **no effect**<br>***IA task*age_quad***<br>$\mathbf{F_{(1,67)}}$**=3.869, *p*=.053** | no effect | no effect | no effect |
| delivery | no effect | **no effect** | **no effec**t<br>*IA task*diagnosis*<br>$F_{(1,68)}$=3.924, *p*=.052 | **right middle frontal gyrus:**<br>$\mathbf{F_{(1,64)}}$**=30.89, $\mathbf{p_{FWE}}$=.011**<br>**left middle frontal gyrus:** | no effect | $\mathbf{F_{(1,68)}}$**=4.70, *p*=.034** | **no effect** | no effect | no effect | no effect | no effect | no effect |

| | | | | |
|---|---|---|---|---|
| | | | | **$F_{(1,64)}$=28.91, $p_{FWE}$=.019** |

Table provides test statistic of region of interest (ROI) analysis and of significant peak voxel(s) for whole-brain analysis. Voxel-level statistics were family-wise error (FWE) corrected for the number of voxels across the whole brain for each test. Significance was defined as pFWE<.05 with a cluster threshold of k≥5. To correct for investigating left and right VS activity separately in the ROI analysis, critical alpha was adjusted to p<.025 based on the Bonferroni procedure. Deviations from results reported in the main text are highlighted in bold. Significant whole-brain results are localized in MNI space and labeled according to the automated anatomical labeling atlas (aal). Abbreviations: TD typically developing, ASD autism spectrum disorder, VS ventral striatum, SID social incentive delay task, MID monetary incentive delay task.

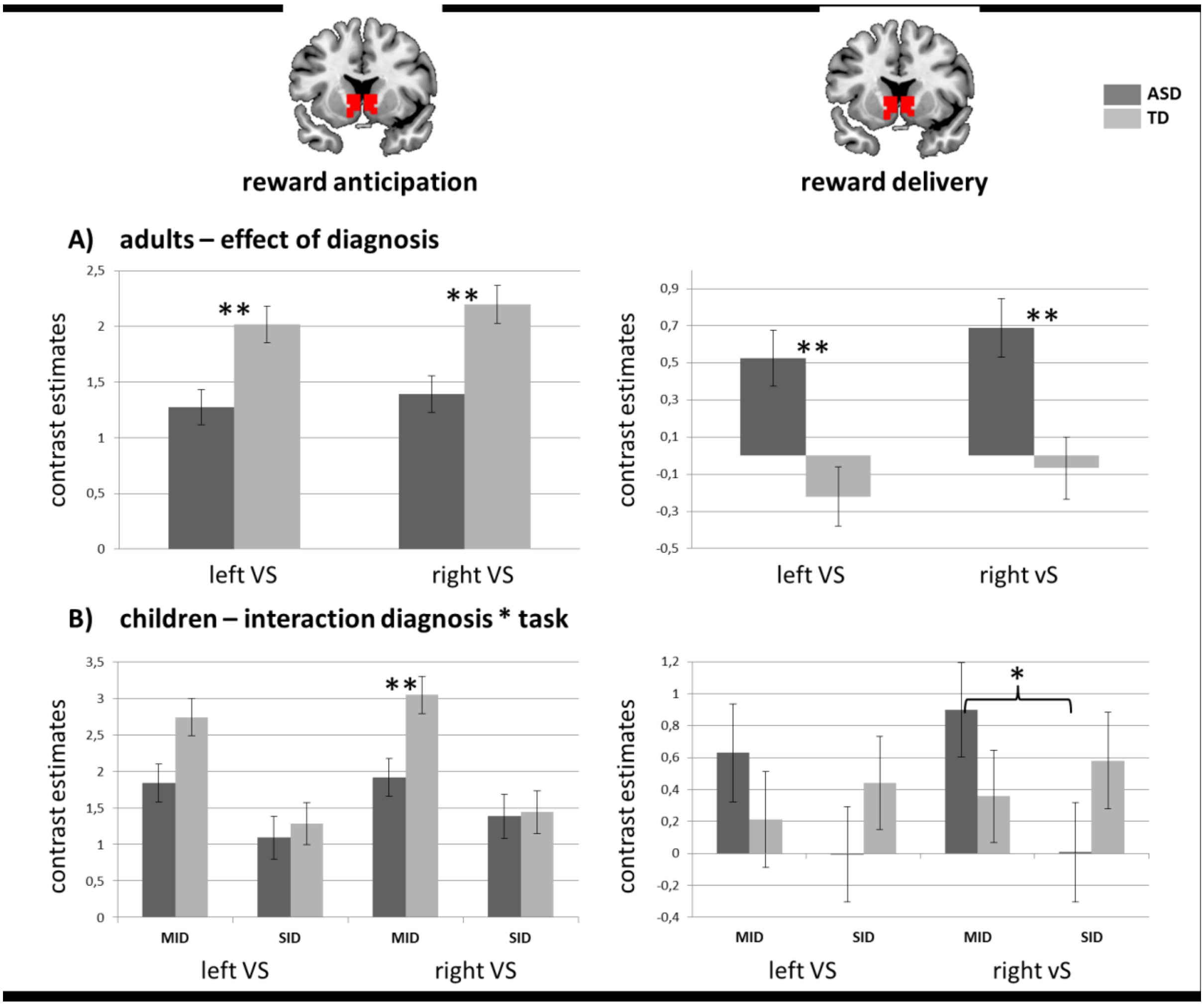

**Figure S4**: Contrast estimates for ventral striatal activation during reward anticipation and delivery for A) adults and B) children. ***p*<.01, **p*<.05.

### ADHD comorbidity

Given the significant comorbidity between ASD and ADHD, and based on the extensive literature on ventral striatal abnormality in ADHD, we investigated the effect of diagnosis while controlling for ADHD comorbidity. Note that information on the presence of a confirmed diagnosis of ADHD was not available in our sample. As a proxy, we calculated an artificial diagnosis applying DSM-V criteria based on symptom scores in the parent- and self-rated ADHD rating scale (33). The pattern of results in the left and right VS was similar when controlling for ADHD comorbidity. The whole-brain level finding of reduced right VS activity in ASD was not replicated when controlling for ADHD, which might, however, also be due to the reduced sample size of individuals with completed ADHD rating-scale.

**Table S5**: Overview over effects of diagnosis and ADHD comorbidity on functional brain activation.

| | TD vs. ASD | | |
|---|---|---|---|
| | whole-brain | left VS | right VS |
| original results (reported in main text) | | | |
| anticipation | *Right VS* $F_{(1,384)}$=22.84, $p_{FWE}$=.017 | $F_{(1,384)}$=14.163, $p$<.001 | $F_{(1,384)}$=18.693, $p$<.001 |
| delivery | no effect | $F_{(1,370)}$=4.829, $p$=.029 | $F_{(1,370)}$=4.719, $p$=.030 |
| results controlling for ADHD comorbidity | | | |
| anticipation | **no effect** | $F_{(1,317)}$=9.895 $p$=.002 | $F_{(1,317)}$=13.187, $p$<=.001 |
| delivery | no effect | $F_{(1,307)}$=7.598 $p$=.006 | $F_{(1,307)}$=5.074 $p$=.025 |

Table provides test statistic of region of interest (ROI) analysis and of significant peak voxel(s) for whole-brain analysis. Voxel-level statistics were family-wise error (FWE) corrected for the number of voxels across the whole brain for each test. Significance was defined as pFWE<.05 with a cluster threshold of k≥5. To correct for investigating left and right VS activity separately in the ROI analysis, critical alpha was adjusted to p<.025 based on the Bonferroni procedure. Deviations from results reported in the main text are highlighted in bold. Significant whole-brain results are localized in MNI space and labeled according to the automated anatomical labeling atlas (aal). Abbreviations: TD typically developing, ASD autism spectrum disorder, ADHD attention-deficit/hyperactivity disorder, VS ventral striatum.

### Site

All analyses were controlled for site. However, to further examine systematic effects while preserving statistical power we employed a leave-one-out approach, recalculating the statistical maps while holding out one site at a time. The pattern of results remained largely the same with vanished whole-brain level effect in the right VS, possibly due to decreased statistical power, when dropping all sites except UCAM. For ROI analysis, the pattern remained largely stable when dropping all sites except when dropping RUN and KCL for reward delivery. As these two sites contributed the largest groups of participants, this is likely due to decreased statistical power.

**Table S6**: Overview over effects of diagnosis and age on functional brain activation after controlling for site using the leave-one-out approach.

| | TD vs. ASD | | | effect of SRS-2 | | |
|---|---|---|---|---|---|---|
| | whole-brain | left VS | right VS | whole-brain | left VS | right VS |
| original results (reported in main text) | | | | | | |
| anticipation | *Right VS* $F_{(1,384)}$=22.84, $p_{FWE}$=.017 | $F_{(1,384)}$=14.163, $p$<.001 | $F_{(1,384)}$=18.693, $p$<.001 | no effect | no effect | no effect |
| delivery | no effect | $F_{(1,370)}$=4.829, $p$=.029 | $F_{(1,370)}$=4.719, $p$=.030 | no effect | no effect | no effect |
| drop site RUNMC | | | | | | |
| anticipation | **no effect** | $F_{(1,270)}$=11.561, $p$=.001 | $F_{(1,270)}$=16.557, $p_{F}$<.001 | no effect | no effect | no effect |
| delivery | no effect | **no effect** | **$F_{(1,257)}$=3.561, $p$=.060** | no effect | no effect | no effect |
| Drop site UMCU | | | | | | |
| anticipation | **no effect** | $F_{(1,315)}$=9.458, $p$=.002 | $F_{(1,315)}$=13.065, $p$<.001 | no effect | no effect | no effect |

| | | | | | | |
|---|---|---|---|---|---|---|
| delivery | no effect | $F_{(1,313)}$=4.777, *p*=.030 | $F_{(1,313)}$=3.929, *p*=.048 | no effect | no effect | no effect |
| Drop site KCL | | | | | | |
| anticipation | **no effect** | $F_{(1,302)}$=11.094, *p*=.001 | $F_{(1,302)}$=13.334, *p*<.001 | no effect | no effect | no effect |
| delivery | no effect | **no effect** | **no effect** | no effect | no effect | no effect |
| Drop site UCAM | | | | | | |
| anticipation | ***left caudate/thalamus*** **$F_{(1,333)}$=22.24, $p_{FWE}$=.025** *right VS* $F_{(1,333)}$=21.97, $p_{FWE}$=.028 | $F_{(1,332)}$=14.269, *p*<.001 | $F_{(1,332)}$=18.623, *p*<.001 | no effect | no effect | no effect |
| delivery | no effect | $F_{(1,318)}$=4.602, *p*=.033 | $F_{(1,318)}$=4.478, *p*=.035 | no effect | no effect | no effect |
| Drop site CIMH | | | | | | |
| anticipation | **no effect** | $F_{(1,340)}$=12.636, *p*<.001 | $F_{(1,340)}$=16.557, *p*<.001 | no effect | no effect | no effect |
| delivery | no effect | $F_{(1,326)}$=5.724, *p*=.017 | $F_{(1,326)}$=5.167, *p*=.024 | no effect | no effect | no effect |
| Drop site UCBM | | | | | | |
| anticipation | **no effect** | $F_{(1,358)}$=12.326, *p*=.001 | $F_{(1,358)}$=16.151, *p*<.001 | no effect | no effect | no effect |
| delivery | | $F_{(1,345)}$=4.049, *p*=.045 | $F_{(1,345)}$=3.860, *p*=.050 | | no effect | no effect |

Table provides test statistic of region of interest (ROI) analysis and of significant peak voxel(s) for whole-brain analysis. Voxel-level statistics were family-wise error (FWE) corrected for the number of voxels across the whole brain for each test. Significance was defined as pFWE<.05 with a cluster threshold of k≥5. To correct for investigating left and right VS activity separately in the ROI analysis, critical alpha was adjusted to p<.025 based on the Bonferroni procedure. Deviations from results reported in the main text are highlighted in bold. Significant whole-brain results are localized in MNI space and labeled according to the automated anatomical labeling atlas (aal). Abbreviations: TD typically developing, ASD autism spectrum disorder, VS ventral striatum, KCL Kings College London, UCBM University Campus Bio-Medico of Rome, UMCU University Medical Centre Utrecht, RUNMC Radboud University Nijmegen Medical Centre, CIMH Central Institute of Mental Health in Mannheim, UCAM University of Cambridge .

## IQ

Level of intellectual abilities was assessed using the Wechsler Abbreviated Scales of Intelligence – Second Edition, WASI-II (Wechsler, 2011) or – in countries where the WASI is not translated (i.e. The Netherlands, Germany and Italy) – the four-subtest short-forms of the German, Dutch or Italian WISC-III/IV (Wechsler, 1991; 2005 for children) or WAIS-III/IV (Wechsler, 1997; Wechsler, 2008 for adults). The shortened versions were used for feasibility reasons to not further prolong the testing sessions for participants. All versions included two verbal subscales (Vocabulary, Similarities) and two non-verbal subscales (Block Design, Matrix Reasoning). To standardise data across sites, IQ was pro-rated from two verbal subtests (vocabulary and similarities) and two performance subtests (matrix reasoning and block design) using an algorithm developed by Sattler (1992) that produces an estimated IQ score that is highly correlated (r = .93) with a Full-Scale IQ obtained by administering the complete test. Age-appropriate national population norms were available for each participating site and these were used to derive standardised estimates of an individual's intellectual functioning. Where recent IQ scores from previous assessments were available (less than 12 months in children; less than 18 months in adolescents and adults) IQ tests were not repeated.

To assess the influence of intellectual ability, full IQ scores were added as additional covariate of no interest to the second level models assessing brain activation differences between individuals with ASD and TD and autism trait scores.

Results were not significantly impacted by controlling for IQ.

**Table S7**: Overview over effects of diagnosis on functional brain activation after controlling for intellectual ability.

| | TD vs. ASD | | | Effect of SRS-2 | | |
|---|---|---|---|---|---|---|
| | whole-brain | left VS | right VS | whole-brain | left VS | right VS |
| original results (reported in main text) | | | | | | |
| anticipation | *right VS* $F_{(1,384)}$=22.84, $p_{FWE}$=.017 | $F_{(1,384)}$=14.163, *p*<.001 | $F_{(1,384)}$=18.693, *p*<.001 | no effect | no effect | no effect |
| delivery | no effect | $F_{(1,370)}$=4.829, *p*=.029 | $F_{(1,370)}$=4.719, *p*=.030 | no effect | no effect | no effect |
| results controlling for IQ | | | | | | |
| anticipation | *Right VS* $F_{(1,381)}$=22.14, $p_{FWE}$=.023 | $F_{(1,383)}$=14.346, *p*<.001 | $F_{(1,383)}$=18.848, *p*)<.001 | no effect | no effect | no effect |
| delivery | no effect | $F_{(1,369)}$=4.774, *p*=.030 | $F_{(1,369)}$=4.684, *p*=.031 | no effect | no effect | no effect |

Table provides test statistic of region of interest (ROI) analysis and of significant peak voxel(s) for whole-brain analysis. Voxel-level statistics were family-wise error (FWE) corrected for the number of voxels across the whole brain for each test. Significance was defined as pFWE<.05 with a cluster threshold of k≥5. To correct for investigating left and right VS activity separately in the ROI analysis, critical alpha was adjusted to p<.025 based on the Bonferroni procedure. Deviations from results reported in the main text are highlighted in bold. Significant whole-brain results are localized in MNI space and labeled according to the automated anatomical labeling atlas (aal). Abbreviations: TD typically developing, ASD autism spectrum disorder, VS ventral striatum.

### Medication

Medication use was confirmed in 82 individuals with ASD and 12 control participants. Most frequently used were psychostimulants and other drugs used to treat ADHD (44.6%), hypnotics/sedatives (31.9%), and antidepressants (24.5%).

Adding a dichotomous covariate indicating medication use (yes/no) to the second level models assessing brain activation did not change the pattern of diagnosis effects on the whole-brain level or within the VS. However, assessing differences between ASD and TD in unmedicated and medicated participants separately yielded no significant whole-brain level and ROI effects, besides a trend for a diagnosis effect in the left VS during reward delivery. This possibly reflects decreased statistical power due to the significantly reduced sample sizes.

**Table S8**: Overview over effects of diagnosis and age on functional brain activation after controlling for site using the leave-one-out approach.

| | TD vs. ASD in unmedicated participants only (n=136) | | | TD vs. ASD in medicated participants only (n=94) | | | TD vs. ASD controlled for effect of medication (whole sample) | | |
|---|---|---|---|---|---|---|---|---|---|
| | whole-brain | left VS | right VS | whole-brain | left VS | right VS | whole-brain | left VS | right VS |
| anticipation | **no effect** | **no effect** | **no effect** | **no effect** | **no effect** | **no effect** | *Right VS* $F_{(1,381)}$=22.14, $p_{FWE}$=.023 | $F_{(1,383)}$=14.302, *p*<.001 | $F_{(1,383)}$=18.525, *p*<.001 |
| delivery | no effect | $F_{(1,123)}$=3.564, *p*=.061 | **no effect** | no effect | **no effect** | **no effect** | no effect | $F_{(1,369)}$=4.596, *p*=.033 | $F_{(1,369)}$=4.433, *p*=.036 |

Table provides test statistic of region of interest (ROI) analysis and of significant peak voxel(s) for whole-brain analysis. Voxel-level statistics were family-wise error (FWE) corrected for the number of voxels across the whole brain for each test. Significance was defined as pFWE<.05 with a cluster threshold of k≥5. To correct for investigating left and right VS activity separately in the ROI analysis, critical alpha was adjusted to p<.025 based on the Bonferroni procedure. Deviations from results reported in the main text are highlighted in bold. Significant whole-brain results are localized in MNI space and labeled according to the automated anatomical labeling atlas (aal). Abbreviations: TD typically developing, ASD autism spectrum disorder, VS ventral striatum.

### Motion

In order to control for potential effects of head motion on our results, we included mean FD as covariate of no interest in the second level models assessing brain activation. Results were not significantly impacted by controlling for head motion. This suggests that the effect of diagnosis is not driven by the fact that individuals with ASD show partly more movement than TD individuals, for example in the SID (see table 1, main text).

**Table S9**: Overview over effects of diagnosis on functional brain activation after controlling for head motion

| | TD vs. ASD | | | effect of SRS-2 | | |
|---|---|---|---|---|---|---|
| | whole-brain | left VS | right VS | whole-brain | left VS | right VS |
| original results (reported in main text) | | | | | | |

| | | | | | | |
|---|---|---|---|---|---|---|
| anticipation | *right VS* $F_{(1,384)}$=22.84, $p_{FWE}$=.017 | $F_{(1,384)}$=14.163, *p*<.001 | $F_{(1,384)}$=18.693, *p*<.001 | no effect | no effect | no effect |
| delivery | no effect | $F_{(1,370)}$=4.829, *p*=.029 | $F_{(1,370)}$=4.719, *p*=.030 | no effect | no effect | no effect |
| Controlling for head motion | | | | | | |
| anticipation | *Right VS:* $F_{(1,383)}$=21.75, $p_{FWE}$=.027 | $F_{(1,382)}$=13.630, *p*<.001 | $F_{(1,382)}$=18.066, *p*<.001 | no effect | no effect | no effect |
| delivery | no effect | $F_{(1,368)}$=6.762, *p*=.010 | $F_{(1,368)}$=6.221, *p*=.013 | no effect | no effect | no effect |

Table provides test statistic of region of interest (ROI) analysis and of significant peak voxel(s) for whole-brain analysis. Voxel-level statistics were family-wise error (FWE) corrected for the number of voxels across the whole brain for each test. Significance was defined as pFWE<.05 with a cluster threshold of k≥5. To correct for investigating left and right VS activity separately in the ROI analysis, critical alpha was adjusted to p<.025 based on the Bonferroni procedure. Deviations from results reported in the main text are highlighted in bold. Significant whole-brain results are localized in MNI space and labeled according to the automated anatomical labeling atlas (aal). Abbreviations: TD typically developing, ASD autism spectrum disorder, VS ventral striatum.

### Sex

All analyses were controlled for sex. However, we additionally explored (1) whether ASD diagnosis effects interacted with sex and (2) if we could replicate our findings in males and females separately.

We found no significant interaction between sex and diagnosis at the whole-brain level or for the right VS. In the left VS, a trend-level significant interaction between diagnosis and sex emerged for reward delivery. Post hoc Bonferroni corrected pairwise comparisons yielded increased left VS activity during reward delivery only in males with ASD compared to male TD (*p*=.004) while in females there was no significant difference between ASD and TD (*p*=.709). Similarly, we could only replicate the effect of diagnosis during reward delivery in male participants (n=272) but not female participants (n=121). This finding likely suggests a lack of statistical power in the female sample, but warrants further inspection in future studies.

**Table S10**: Overview over effects of diagnosis and age on functional brain activation in male and female participants separately.

| | TD vs. ASD | | | effect of SRS-2 | | |
|---|---|---|---|---|---|---|
| | whole-brain | left VS | right VS | whole-brain | left VS | right VS |
| sex*diagnosis interaction | | | | | | |
| anticipation | no effect | no effect | no effect | | no effect | no effect |
| delivery | no effect | $F_{(1,369)}$=3.885, *p*=.049 | no effect | | no effect | no effect |
| original results (reported in main text) | | | | | | |
| anticipation | *right VS* $F_{(1,384)}$=22.84, | $F_{(1,384)}$=14.163, *p*<.001 | $F_{(1,384)}$=18.693, *p*<.001 | no effect | no effect | no effect |

| | | | | | | |
|---|---|---|---|---|---|---|
| | $p_{FWE}$=.017 | | | | | |
| delivery | no effect | $F_{(1,370)}$=4.829, $p$=.029 | $F_{(1,370)}$=4.719, $p$=.030 | no effect | no effect | no effect |
| only male participants (n=272) | | | | | | |
| anticipation | no effect | $F_{(1,264)}$=9.017, $p$=.003 | $F_{(1,264)}$=13.509 , $p$<.001 | no effect | no effect | no effect |
| delivery | no effect | $F_{(1,252)}$=8.632, $p$=.004 | $F_{(1,252)}$=7.854, $p$=.005 | no effect | no effect | no effect |
| only female participants (n=121) | | | | | | |
| anticipation | no effect | $F_{(1,113)}$=6.352, $p$=.013 | $F_{(1,113)}$=6.589, $p$=.012 | no effect | no effect | no effect |
| delivery | no effect | **no effect** | **no effect** | no effect | no effect | no effect |

Table provides test statistic of region of interest (ROI) analysis and of significant peak voxel(s) for whole-brain analysis. Voxel-level statistics were family-wise error (FWE) corrected for the number of voxels across the whole brain for each test. Significance was defined as pFWE<.05 with a cluster threshold of k≥5. To correct for investigating left and right VS activity separately in the ROI analysis, critical alpha was adjusted to p<.025 based on the Bonferroni procedure. Deviations from results reported in the main text are highlighted in bold. Significant whole-brain results are localized in MNI space and labeled according to the automated anatomical labeling atlas (aal). Abbreviations: TD typically developing, ASD autism spectrum disorder, VS ventral striatum.

### Handedness

The majority (68.9%, n=271) of participants were right handed. However, to rule out any impact of handedness on brain activation we repeated the main analyses in righthanded participants only, replicating group differences during reward anticipation and delivery.

**Table S11**: Overview over effects of diagnosis and age on functional brain activation in righthanders (n=271).

| | TD vs. ASD | | | effect of SRS-2 | | |
|---|---|---|---|---|---|---|
| | whole-brain | left VS | right VS | whole-brain | left VS | right VS |
| anticipation | **no effect** | $F_{(1,263)}$=10.161, $p$=.002 | $F_{(1,263)}$=12.159, $p$=.001 | no effect | no effect | no effect |
| delivery | no effect | $F_{(1,254)}$=4.529, $p$=.034 | $F_{(1,254)}$=5.511, $p$=.020 | no effect | no effect | no effect |

Table provides test statistic of region of interest (ROI) analysis and of significant peak voxel(s) for whole-brain analysis. Voxel-level statistics were family-wise error (FWE) corrected for the number of voxels across the whole brain for each test. Significance was defined as pFWE<.05 with a cluster threshold of k≥5. To correct for investigating left and right VS activity separately in the ROI analysis, critical alpha was adjusted to p<.025 based on the Bonferroni procedure. Deviations from results reported in the main text are highlighted in bold. Significant whole-brain results are localized in MNI space and labeled according to the automated anatomical labeling atlas (aal). Abbreviations: TD typically developing, ASD autism spectrum disorder, VS ventral striatum.

1. Loth E, Charman T, Mason L, Tillmann J, Jones EJH, Wooldridge C, et al. (2017): The EU-AIMS Longitudinal European Autism Project (LEAP): design and methodologies to identify and validate stratification biomarkers for autism spectrum disorders. *Molecular autism*. 8:24.

2. Moessnang C, Schafer A, Bilek E, Roux P, Otto K, Baumeister S, et al. (2016): Specificity, reliability and sensitivity of social brain responses during spontaneous mentalizing. *Social cognitive and affective neuroscience*.
3. Plichta MM, Grimm O, Morgen K, Mier D, Sauer C, Haddad L, et al. (2014): Amygdala habituation: a reliable fMRI phenotype. *NeuroImage*. 103:383-390.
4. Plichta MM, Schwarz AJ, Grimm O, Morgen K, Mier D, Haddad L, et al. (2012): Test-retest reliability of evoked BOLD signals from a cognitive-emotive fMRI test battery. *NeuroImage*. 60:1746-1758.
5. Jenkinson M, Bannister P, Brady M, Smith S (2002): Improved optimization for the robust and accurate linear registration and motion correction of brain images. *NeuroImage*. 17:825-841.
6. Power JD, Barnes KA, Snyder AZ, Schlaggar BL, Petersen SE (2012): Spurious but systematic correlations in functional connectivity MRI networks arise from subject motion. *NeuroImage*. 59:2142-2154.
7. Neuhaus E, Bernier RA, Beauchaine TP (2015): Electrodermal Response to Reward and Non-Reward Among Children With Autism. *Autism research : official journal of the International Society for Autism Research*. 8:357-370.
8. Demurie E, Roeyers H, Baeyens D, Sonuga-Barke E (2011): Common alterations in sensitivity to type but not amount of reward in ADHD and autism spectrum disorders. *Journal of child psychology and psychiatry, and allied disciplines*. 52:1164-1173.
9. Kohls G, Antezana L, Mosner MG, Schultz RT, Yerys BE (2018): Altered reward system reactivity for personalized circumscribed interests in autism. *Molecular autism*. 9:9.
10. Kohls G, Schulte-Ruther M, Nehrkorn B, Muller K, Fink GR, Kamp-Becker I, et al. (2013): Reward system dysfunction in autism spectrum disorders. *Social cognitive and affective neuroscience*. 8:565-572.
11. Kohls G, Thonessen H, Bartley GK, Grossheinrich N, Fink GR, Herpertz-Dahlmann B, et al. (2014): Differentiating neural reward responsiveness in autism versus ADHD. *Developmental cognitive neuroscience*. 10:104-116.
12. van Dongen EV, von Rhein D, O'Dwyer L, Franke B, Hartman CA, Heslenfeld DJ, et al. (2015): Distinct effects of ASD and ADHD symptoms on reward anticipation in participants with ADHD, their unaffected siblings and healthy controls: a cross-sectional study. *Molecular autism*. 6:48.
13. Greene RK, Spanos M, Alderman C, Walsh E, Bizzell J, Mosner MG, et al. (2018): The effects of intranasal oxytocin on reward circuitry responses in children with autism spectrum disorder. *Journal of neurodevelopmental disorders*. 10:12.
14. Delmonte S, Balsters JH, McGrath J, Fitzgerald J, Brennan S, Fagan AJ, et al. (2012): Social and monetary reward processing in autism spectrum disorders. *Molecular autism*. 3:7.
15. Galvan A (2010): Adolescent development of the reward system. *Frontiers in human neuroscience*. 4:6.
16. Ernst M, Daniele T, Frantz K (2011): New perspectives on adolescent motivated behavior: attention and conditioning. *Developmental cognitive neuroscience*. 1:377-389.
17. Bjork JM, Knutson B, Fong GW, Caggiano DM, Bennett SM, Hommer DW (2004): Incentive-elicited brain activation in adolescents: similarities and differences from young adults. *J Neurosci*. 24:1793-1802.
18. Bjork JM, Smith AR, Chen G, Hommer DW (2010): Adolescents, adults and rewards: comparing motivational neurocircuitry recruitment using fMRI. *PloS one*. 5:e11440.
19. Geier CF, Terwilliger R, Teslovich T, Velanova K, Luna B (2010): Immaturities in reward processing and its influence on inhibitory control in adolescence. *Cerebral cortex*. 20:1613-1629.
20. May JC, Delgado MR, Dahl RE, Stenger VA, Ryan ND, Fiez JA, et al. (2004): Event-related functional magnetic resonance imaging of reward-related brain circuitry in children and adolescents. *Biological psychiatry*. 55:359-366.
21. Van Leijenhorst L, Gunther Moor B, Op de Macks ZA, Rombouts SA, Westenberg PM, Crone EA (2010): Adolescent risky decision-making: neurocognitive development of reward and control regions. *NeuroImage*. 51:345-355.
22. Galvan A, Hare TA, Parra CE, Penn J, Voss H, Glover G, et al. (2006): Earlier development of the accumbens relative to orbitofrontal cortex might underlie risk-taking behavior in adolescents. *J Neurosci*. 26:6885-6892.
23. Ernst M, Nelson EE, Jazbec S, McClure EB, Monk CS, Leibenluft E, et al. (2005): Amygdala and nucleus accumbens in responses to receipt and omission of gains in adults and adolescents. *NeuroImage*. 25:1279-1291.
24. Schreuders E, Braams BR, Blankenstein NE, Peper JS, Guroglu B, Crone EA (2018): Contributions of Reward Sensitivity to Ventral Striatum Activity Across Adolescence and Early Adulthood. *Child Dev*. 89:797-810.

25. Lamm C, Benson BE, Guyer AE, Perez-Edgar K, Fox NA, Pine DS, et al. (2014): Longitudinal study of striatal activation to reward and loss anticipation from mid-adolescence into late adolescence/early adulthood. *Brain and cognition*. 89:51-60.
26. Hoogendam JM, Kahn RS, Hillegers MH, van Buuren M, Vink M (2013): Different developmental trajectories for anticipation and receipt of reward during adolescence. *Developmental cognitive neuroscience*. 6:113-124.
27. Smith AB, Halari R, Giampetro V, Brammer M, Rubia K (2011): Developmental effects of reward on sustained attention networks. *NeuroImage*. 56:1693-1704.
28. Duverne S, Koechlin E (2017): Rewards and Cognitive Control in the Human Prefrontal Cortex. *Cerebral cortex*. 27:5024-5039.
29. Kumar P, Pisoni A, Bondy E, Kremens R, Singleton P, Pizzagalli DA, et al. (2019): Delineating the Social Valuation Network in Adolescents. *Social cognitive and affective neuroscience*.
30. Ruff CC, Fehr E (2014): The neurobiology of rewards and values in social decision making. *Nat Rev Neurosci*. 15:549-562.
31. Dreher JC, Meyer-Lindenberg A, Kohn P, Berman KF (2008): Age-related changes in midbrain dopaminergic regulation of the human reward system. *Proc Natl Acad Sci U S A*. 105:15106-15111.
32. Dhingra I, Zhang S, Zhornitsky S, Le TM, Wang W, Chao HH, et al. (2020): The effects of age on reward magnitude processing in the monetary incentive delay task. *NeuroImage*. 207:116368.
33. DuPaul GJ, Power TJ, Anastopoulos AD, Reid R (1998): *ADHD Rating Scale—IV: Checklists, norms, and clinical interpretation*. New York, NY, US: Guilford Press.
